# Sex differences in the impact of electronic nicotine vapor on corticotropin-releasing factor receptor 1 neurons in the mouse ventral tegmental area

**DOI:** 10.1101/2022.08.19.504534

**Authors:** ManHua Zhu, Neil G Rogers, Jasmine V Jahad, Melissa A Herman

## Abstract

Nicotine engages dopamine neurons in the ventral tegmental area (VTA) to encode reward and drive the development of nicotine addiction, however how nicotine selectively alters other VTA populations remains to be determined. Here, we used male and female CRF1-GFP mice and nicotine vapor exposure to examine the effects of nicotine in VTA corticotropin-releasing factor receptor 1 (CRF1) neurons. We use immunohistochemistry and electrophysiology to examine neuronal activity, excitability, and inhibitory signaling. We found that VTA CRF1 neurons are mainly dopaminergic and project to the nucleus accumbens (^VTA-NAc^CRF1 neurons). ^VTA-NAc^CRF1 neurons show greater phasic inhibition in naïve females and greater focal nicotine-induced increases in firing in naïve males. Following acute nicotine vapor exposure, phasic inhibition was not altered, but focal nicotine-induced tonic inhibition was enhanced in females and diminished in males. Acute nicotine vapor exposure did not affect firing in ^VTA-NAc^CRF1 neurons, but females showed lower baseline firing and higher focal nicotine-induced firing. Activity (cFos) was increased in the CRF1 dopaminergic VTA population in both sexes, but with greater increases in females. Following chronic nicotine vapor exposure, both sexes displayed reduced basal phasic inhibition and the sex difference in tonic inhibition following acute vapor exposure was no longer observed. Additionally, activity of the CRF1 dopaminergic VTA population was no longer elevated in either sex. These findings reveal sex- and exposure-dependent changes in mesolimbic VTA-NAc CRF1 neuronal activity, inhibitory signaling, and nicotine sensitivity following nicotine vapor exposure. These changes potentially contribute to nicotine-dependent behaviors and the intersection between stress, anxiety, and addiction.

## Introduction

Canonically, nicotine’s addictive properties result from activation of nicotinic acetylcholine receptors on dopaminergic neurons in the ventral tegmental area (VTA) driving dopamine release in the nucleus accumbens (NAc). Nicotine produces distinct temporal patterns of excitatory glutamatergic and inhibitory GABAergic inputs onto VTA dopamine neurons driving overall increases in excitability^1, 2^. Although nicotine typically increases VTA dopamine neuron activity, nicotine is also inhibitory in amygdala-projecting VTA dopamine neurons and increases anxiety-like behaviors^3^. *In vivo* behavioral studies have found that rodents will self-administer nicotine intravenously^4–7^, intracranially into the VTA^8–10^, and through inhalation^11, 12^. However, more studies are needed to examine how nicotine vapor delivered through electronic nicotine delivery systems (ENDS) impacts specific VTA neuronal populations and mesolimbic reward circuit activity.

Nicotine also plays an important role in modulating stress and anxiety behaviors and stress-induced relapse to drug seeking^13–15^. Human studies report that women are more likely to smoke to manage stress and experience anxiety more intensely during abstinence^16^. The corticotropin-releasing factor (CRF) system regulates stress responding and has been implicated in cocaine^17–19^, alcohol^20–23^, and nicotine addiction^24, 25^. Nicotine activates the hypothalamic-pituitary-adrenal (HPA) axis to release CRF in the thalamus^26^. Chronic exposure to nicotine alters basal HPA axis activity^26^ and increases CRF mRNA expression in the VTA^24^. CRF peptide increases VTA dopamine neuron firing via CRF receptor 1 (CRF1)^27, 28^. Previous studies found that VTA CRF1 neurons mainly project to the NAc core and activation of these neurons’ cell bodies and terminals coordinate reward reinforcement behavior and enhance dopamine release, respectively^29^. Conversely, knockout of CRF1 in midbrain dopamine neurons has been shown to increase anxiety-like behavior^30^ and CRF1 antagonism reduces footshock-induced reinstatement of nicotine seeking^31^.

Nicotine’s effects in the VTA and modulation of anxiety and stress have been well-studied, however, less is known about CRF1 VTA neurons and their response to electronic nicotine vapor, especially in females. The emergence of ENDS products provides an opportunity for pre-clinical studies to examine sex- and population-specific sensitivity to nicotine vapor exposure. Here, we used male and female CRF1 transgenic mice to determine how acute and chronic nicotine vapor exposure dysregulates activity and inhibitory control of VTA CRF1 neurons.

## Results

### Immunohistochemical characterization of VTA CRF1 neurons

We observed broad distribution of GFP+ (CRF1) neurons across the VTA (**Fig 1A top**) and colocalization of CRF1 and tyrosine hydroxylase (TH+), a marker of dopaminergic neurons (**Fig 1A bottom**). Within TH-expressing VTA neurons, ~40% co-expressed CRF1 (**Fig 1B**). Within CRF1-expressing VTA neurons, ~80% co-expressed TH (**Fig 1C**). No sex differences were observed in co-expression of either CRF1 or TH populations (**Fig 1 B, C**). To investigate whether CRF1 neurons project to the nucleus accumbens (NAc), we injected AAV-hSyn-DIO-eGFP into the VTA (**Fig 1D**) of CRF1-cre mice and observed terminal expression in the NAc (**Fig 1E**). These data indicate that CRF1 neurons in the VTA are predominantly dopaminergic and are part of the mesolimbic VTA-NAc circuitry.

**Fig 1.**
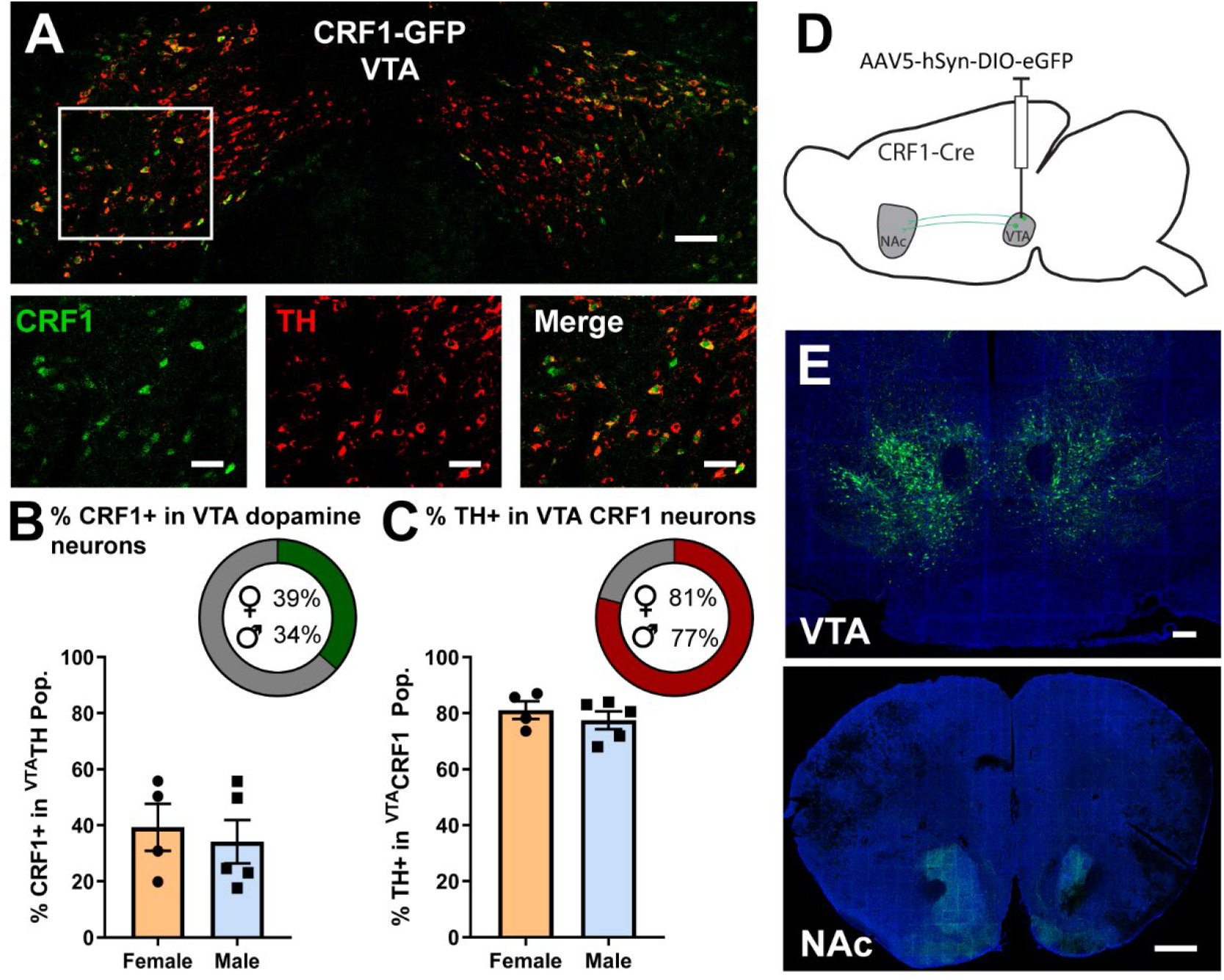
Immunohistochemical characterization of VTA CRF1 neurons. **(A)** Top: Representative image of a coronal section of VTA, scale bar=100 μm. Bottom: Magnified view (white box in top image) of CRF1 (GFP), TH, and merged channels, scale bar=50 μm. **(B)** Percentage of TH neurons in the VTA that are CRF1+ in females (N=4) and males (N=5). Inset: averaged percentages in female and male mice. **(C)** Percentage of CRF1 neurons in the VTA that are TH+ in females (N=4) and males (N=5). Inset: averaged percentages in female and male mice. **(D)** Schematic of viral strategy to probe VTA CRF1 neuron projections. **(E)** Top: VTA injection site, scale bar=200 μm. Bottom: Terminal expression in the NAc, scale bar=500 μm. Data represented as mean ± SEM.

### Electrophysiological characterization of VTA CRF1 neurons

To determine electrophysiological characteristics of VTA CRF1 neurons that project to NAc (^**VTA-NAc**^**CRF1 neurons**), we injected retrograde beads into the NAc of CRF1-GFP mice (**Fig 2A**) and performed slice electrophysiology in the VTA (**Fig 2B left**). Neurons co-expressing GFP (CRF1) and beads were targeted for recording (**Fig 2B inset**). We found no significant differences in membrane properties, except membrane resistance was higher in males than in females (*p=0.0170, t=2.505, unpaired t-test with Welch’s correction; **Fig 2C**). Voltage-clamp recordings of spontaneous inhibitory post-synaptic currents (sIPSCs, **Fig 2D**) were performed and sIPSC frequency was significantly higher in females than males (*p=0.0331, t=2.375, unpaired t-test with Welch’s correction; **Fig 2E**) but sIPSC amplitude showed no sex differences (**Fig 2F**). Focal nicotine (1 μM) did not alter sIPSC frequency (**Fig 2G, H**) or amplitude (**Fig 2I, J**) in either sex, however females showed overall higher sIPSC frequency than males (main effect of sex *p=0.0218, F (1, 20) = 6.185), 2-way ANOVA; **Fig 2G**). In contrast, focal nicotine induced a tonic inhibitory current in ^VTA-NAc^CRF1 neurons of both sexes (**Fig 2K, L**) that was partially reversed by 100 μM gabazine (**Fig 2M**). Cell-attached recordings of spontaneous firing (**Fig 2N**) were performed and we found no sex differences in baseline firing (**Fig 2O**) or nicotine-induced firing (**Fig 2P**). However, focal nicotine induced a significant (~20%) increase in normalized firing frequency in males (*p=0.0271, t=2.517, one sample t-test; **Fig 2Q**). These data indicate that ^VTA-NAc^CRF1 neurons display greater baseline phasic inhibition in naïve females and greater focal nicotine-induced firing in naïve males. Focal nicotine does not alter phasic inhibition but induces tonic inhibition in both sexes.

**Fig 2.**
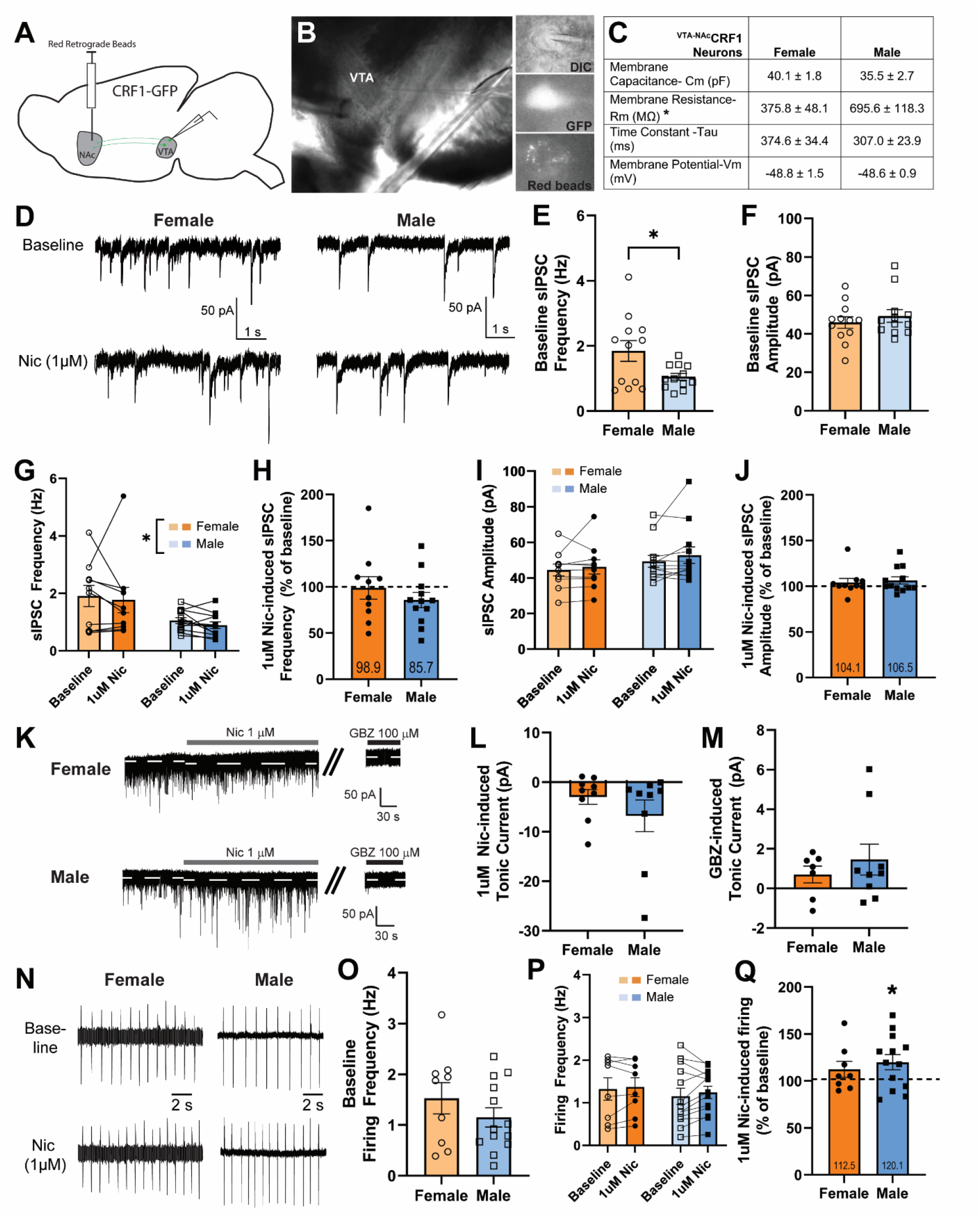
Electrophysiological properties and sensitivity to focal nicotine in ^VTA-NAc^CRF1 neurons. **(A)** Schematic of red retrobead injection in the NAc and electrophysiological recording in the VTA in CRF1-GFP mice. **(B)** Left: Coronal section of VTA (4x) with recording electrode (right) and y-tube (left). Inset: recorded neuron (60x) with differential interference contrast (DIC, top), GFP/CRF1 expression (middle), and red bead expression (bottom). **(C)** Membrane properties of ^VTA-NAc^CRF1 neurons from female (N=17 cells, N=8 mice) and male (N=28 cells, N=11 mice) mice. Males showed higher membrane resistance than females (*p=0.0170, t=2.505, unpaired t-test with Welch’s correction). **(D)** Example traces of spontaneous inhibitory post-synaptic currents (sIPSCs) in ^VTA-NAc^CRF1 neurons from female (left) and male (right) mice at baseline and after 1 μM nicotine application. **(E)** Baseline sIPSC frequency in ^VTA-NAc^CRF1 neurons were higher in female (N=12 cells, N=6 mice) than male (N=12 cells, N=8 mice) mice (*p=0.0331, t=2.375, unpaired t-test with Welch’s correction). **(F)** Baseline sIPSC amplitude in ^VTA-NAc^CRF1 neurons from female (N=12 cells, N=6 mice) and male (N=12 cells, N=8 mice) mice. **(G)** sIPSC frequency at baseline and after 1 μM nicotine application in ^VTA-NAc^CRF1 neurons from female (N=10 cells, N=5 mice) and male (N=12 cells, N=8 mice) mice. Main effect of sex *p=0.0218, F (1, 20) = 6.185), 2-way ANOVA. **(H)** 1 μM nicotine-induced change in sIPSC frequency normalized to baseline in ^VTA-NAc^CRF1 neurons from female (N=10 cells, N=5 mice) and male (N=12 cells, N=8 mice) mice. **(I)** sIPSC amplitude at baseline and after 1 μM nicotine application in ^VTA-NAc^CRF1 neurons from female (N=10 cells, N=5 mice) and male (N=12 cells, N=8 mice) mice. **(J)** 1 μM nicotine-induced change in sIPSC amplitude normalized to baseline in ^VTA-NAc^CRF1 neurons from female (N=10 cells, N=5 mice) and male (N=12 cells, N=8 mice) mice. **(K)** Example traces of 1 μM nicotine-induced and gabazine (GBZ, 100 μM) reversed tonic current in ^VTA-NAc^CRF1 neurons of female (top) and male (bottom) mice. **(L)** 1 μM nicotine-induced tonic current in ^VTA-NAc^CRF1 neurons of female (N=9 cells, N=4 mice) and male (N=9 cells, N=6 mice) mice. **(M)** 100 μM Gabazine (GBZ)-induced reversal of tonic current in ^VTA-NAc^CRF1 neurons of female (N=7 cells, N=4 mice) and male (N=9 cells, N=6 mice) mice. **(N)** Cell-attached firing in ^VTA-NAc^CRF1 neurons from female (left) and male (right) mice at baseline and after 1 μM nicotine application. **(O)** Baseline firing frequency in ^VTA-NAc^CRF1 neurons of female (N=9 cells, N=5 mice) and male (N=13 cells, N=9 mice) mice. **(P)** Firing frequency before and after 1 μM nicotine application in ^VTA-NAc^CRF1 neurons from female (N=8 cells, N=5 mice) and male (N=13 cells, N=9 mice) mice. **(Q)** 1 μM nicotine-induced change in firing frequency normalized to baseline in ^VTA-NAc^CRF1 neurons from female (N=8 cells, N=5 mice) and male (N=13 cells, N=9 mice, *p=0.0271, t=2.517, one sample t-test) mice. Data are represented as mean ± SEM.

### Serum nicotine metabolite levels following acute and chronic electronic nicotine vapor exposure

Acute vapor exposure consisted of a 3-hr session (3-sec vapor deliveries every 10 minutes, **Fig 3A, left**). Chronic exposure consisted of daily 3-hr sessions for 28 days (**Fig 6A, left**). Mice exposed to acute 12% nicotine vapor showed significantly higher serum nicotine (main effect of vapor content ****p<0.0001, F(1,40)=32.75, Female: PG/VG vs Nic ***p=0.0003, Male: PG/VG vs Nic **p=0.0055, 2-way ANOVA post hoc Tukey’s; **Supp Fig 1A, left**) and serum cotinine (main effect of vapor content ****p<0.0001, F(1,40)=53.80, Female: PG/VG vs Nic ****p<0.0001, Male: PG/VG vs Nic ****p<0.0001, 2-way ANOVA post hoc Tukey’s; **Supp Fig 1A, right**) compared to PG/VG controls. Mice exposed to chronic 12% nicotine vapor showed significantly higher serum nicotine (main effect of vapor content ****p<0.0001, F(1,30)=57.68, Female: PG/VG vs Nic ****p<0.0001, Male: PG/VG vs Nic ***p=0.0002, 2-way ANOVA post hoc Tukey’s; **Supp Fig 1B, left**) and serum cotinine (main effect of vapor content ****p<0.0001, F(1,30)=41.95, Female: PG/VG vs Nic **p=0.0077, Male: PG/VG vs Nic ****p<0.0001, 2-way ANOVA post hoc Tukey’s; **Supp Fig 1B, right**) compared to PG/VG controls. Together, these data demonstrate that the delivery of nicotine through electronic vapor is an effective route of nicotine administration that produces significant serum nicotine and cotinine levels in animals following acute and chronic exposure.

**Fig 3.**
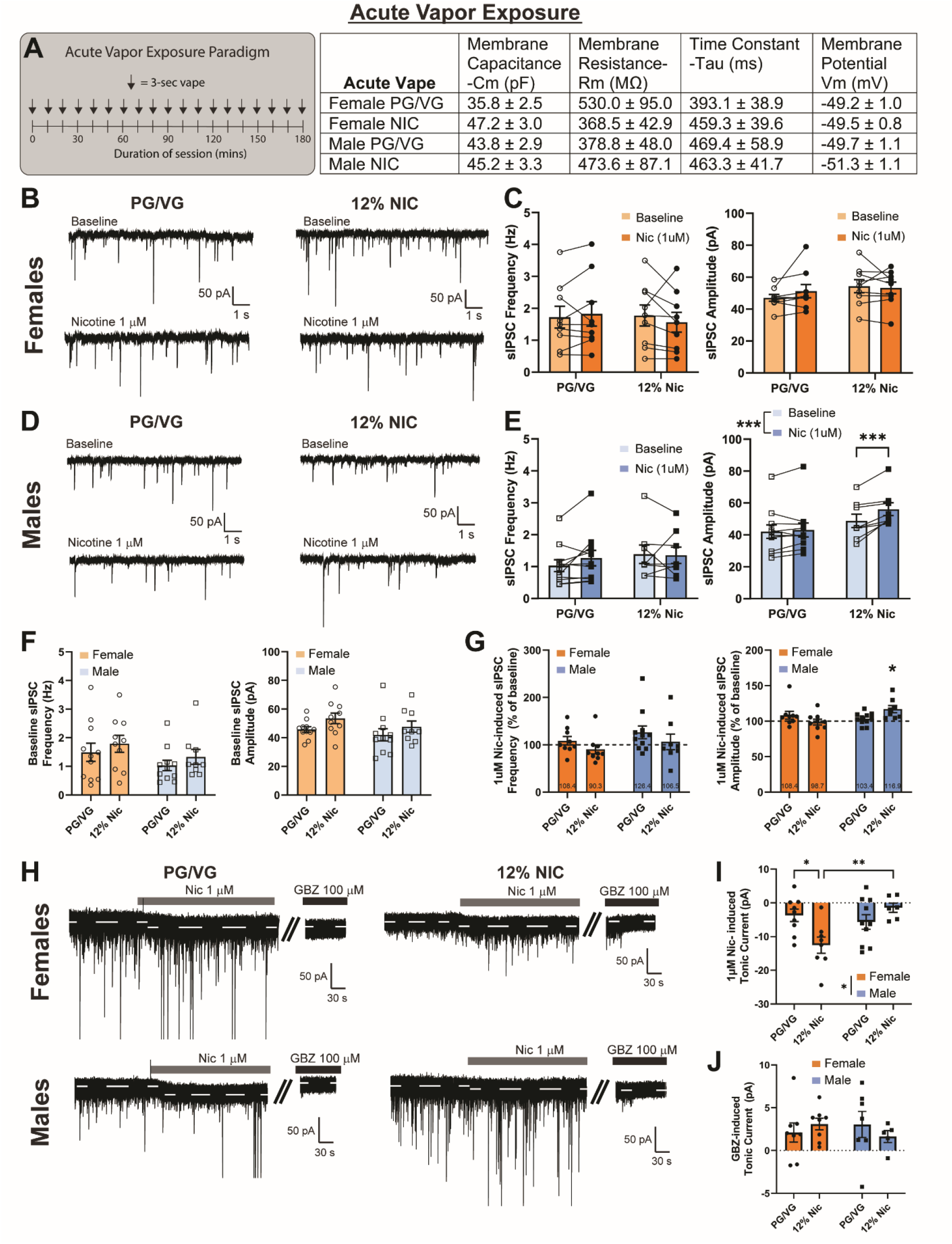
Inhibitory signaling in ^VTA-NAc^CRF1 neurons following acute vapor exposure. **(A)** Acute vapor exposure paradigm (left) and membrane properties (right) of ^VTA-NAc^CRF1 neurons from male and female mice exposed to acute PG/VG (Female N=23 cells, N=6 mice; Male N=22 cells, N=5 mice) or 12% nicotine vapor (Female N=15 cells, N=4 mice; Male N=21 cells, N=5 mice). There’s a main effect of vapor content on membrane capacitance (*p=0.0353, F(1,77)=4.591, 2-way ANOVA). **(B)** Spontaneous inhibitory post-synaptic currents (sIPSCs) in ^VTA-NAc^CRF1 neurons from female mice exposed to PG/VG (left) or 12% nicotine (right) at baseline and after 1 μM nicotine application. **(C)** sIPSC frequency (left) and amplitude (right) in ^VTA-NAc^CRF1 neurons from PG/VG (N=9 cells, N=5 mice) or 12% nicotine (N=9 cells, N=4 mice) exposed female mice at baseline and after 1 μM nicotine application. **(D)** Spontaneous inhibitory post-synaptic currents (sIPSCs) in ^VTA-NAc^CRF1 neurons from male mice exposed to PG/VG (left) or 12% nicotine (right) at baseline and after 1 μM nicotine application. **(E)** sIPSC frequency (left) and amplitude (right) in ^VTA-NAc^CRF1 neurons from PG/VG (N=11 cells, N=4 mice) or 12% nicotine (N=8 cells, N=5 mice) exposed male mice at baseline and after 1 μM nicotine application. sIPSC amplitude showed vapor content x focal nicotine interaction (**p=0.0044, F (1, 17) = 10.77), main effect of focal nicotine (***p=0.0004, F (1, 17) = 19.40), and12% Nic: baseline vs 1uM nic (***p=0.0002), RM 2-way ANOVA post hoc Sidak’s. **(F)** Baseline sIPSC frequency (left) and amplitude (right) in ^VTA-NAc^CRF1 neurons from female and male mice exposed to PG/VG (Female N=11 cells, N=5 mice; Male N= 11 cells, N=4 mice) or 12% nicotine (Female N=10 cells, N=4 mice; Male N= 8 cells, N=5 mice). **(G)** 1 μM nicotine-induced change in sIPSC frequency (left) and amplitude (right) normalized to baseline in ^VTA-NAc^CRF1 neurons from female and male mice exposed to PG/VG (Female N=9 cells, N=5 mice; Male N= 11 cells, N=4 mice) or 12% nicotine (Female N=9 cells, N=4 mice; Male N= 8 cells, N=5 mice). Normalized sIPSC amplitude show vapor content x sex interaction (*p=0.0101, F (1, 33) = 7.454, 2-way ANOVA) and specifically, the male 12% Nic group shown significant increase from baseline (*p=0.0114, t=3.405, one sample t-test). **(H)** Example traces of 1 μM nicotine-induced and gabazine (GBZ) reversed tonic current in ^VTA-NAc^CRF1 neurons from female and male mice exposed to PG/VG (left) or 12% nicotine (right). **(I)** 1 μM nicotine-induced tonic current in ^VTA-NAc^CRF1 neurons from female and male mice exposed to PG/VG (Female N=9 cells, N=5 mice; Male N=10 cells, N=4 mice) or 12% nicotine (Female N=8 cells, N=4 mice; Male N= 6 cells, N=3 mice). Tonic current showed a vapor content x sex interaction (**p=0.0045, F (1, 29) = 9.486), main effect of sex (*p=0.0389, F (1, 29) = 4.682) and post hoc Tukey’s significance in Female: PG/VG vs Nic (*p=0.0240) and Female Nic vs. Male Nic (**p=0.0091) by 2-way ANOVA. **(J)** Gabazine (GBZ, 100 μM) reversal of tonic current in ^VTA-NAc^CRF1 neurons of female and male mice exposed to PG/VG (Female N=8 cells, N=4 mice; Male N=7 cells, N=4 mice) or 12% nicotine (Female N=8 cells, N=4 mice; Male N=6 cells, N=3 mice). Data are represented as mean ± SEM.

### The effects of acute vapor exposure on inhibitory signaling in ^VTA-NAc^CRF1 neurons

We exposed CRF1-GFP mice to acute vapor (**Fig 3A, left**) and performed slice electrophysiology targeting ^VTA-NAc^CRF1 neurons (**Fig 2A-B**). We found no significant differences in membrane properties (**Fig 3A, right**), except membrane capacitance (main effect of vapor content *p=0.0353, F(1,77)=4.591, 2-way ANOVA; p>0.05 by Tukey’s multiple comparisons test). In females (**Fig 3B**) and males (**Fig 3D**), there was no significant difference in baseline sIPSC frequency (**Fig 3F, left**) or amplitude (**Fig 3F, right**). In females, focal nicotine (1 μM) produced no change in sIPSC frequency (**Fig 3C, left and Fig 3G, left**) or amplitude (**Fig 3C, right and Fig 3G, right**). In males, focal nicotine (1 μM) also produced no change in sIPSC frequency (**Fig 3E, left and Fig 3G, left**) but sIPSC amplitude was significantly increased, mainly driven by the 12% nicotine-exposed group (vapor content x focal nicotine interaction **p=0.0044, F (1, 17) = 10.77; main effect of focal nicotine ***p=0.0004, F (1, 17) = 19.40; 12% Nic: baseline vs 1uM nic ***p=0.0002, RM 2-way ANOVA post hoc Sidak’s; **Fig 3E right**). Normalized sIPSC amplitude was significantly increased in 12% nicotine-exposed males (*p=0.0114, t=3.405, one sample t-test; **Fig 3G, right**). Focal nicotine (1 μM) induced a tonic current that was comparable in PG/VG groups but 12% nicotine-exposed females showed enhanced tonic current whereas 12% nicotine-exposed males showed reduced tonic current (vapor content x sex interaction **p=0.0045, F (1, 29) = 9.486; main effect of sex *p=0.0389, F (1, 29) = 4.682; Female: PG/VG vs Nic *p=0.0240, Female Nic vs. Male Nic **p=0.0091, 2-way ANOVA post hoc Tukey’s; **Fig 3H, I**). Gabazine (GBZ, 100 μM) partially reversed the focal nicotine-induced tonic current (**Fig 3J**). These data suggest that phasic inhibition in ^VTA-NAc^CRF1 neurons is largely unaffected but focal nicotine-induced tonic inhibition was enhanced in females and diminished in males following acute nicotine vapor exposure.

### The effects of acute vapor exposure on spontaneous firing of ^VTA-NAc^CRF1 neurons

We performed cell-attached recordings in female (**Fig 4A, B**) and male (**Fig 4C, D**) mice exposed to acute PG/VG or 12% nicotine vapor. We observed higher baseline firing frequency in males as compared to females (main effect of sex *p=0.0282, F (1, 49) = 5.114, 2-way ANOVA; **Fig 4E**), however with a marginal overall effect size. Focal nicotine (1 μM) in females (**Fig 4A, B**) and males (**Fig 4C, D**) exposed to PG/VG or 12% nicotine vapor produced no change in firing frequency. However, when firing frequency was normalized to baseline, females exposed to either PG/VG or 12% nicotine showed significantly higher firing frequency as compared to males (main effect of sex *p=0.0253, F (1, 37) = 5.437, 2-way ANOVA; **Fig 4F**). These data suggest that acute exposure does not affect spontaneous firing in ^VTA-NAc^CRF1 neurons. However, females exposed to either PG/VG or 12% nicotine vapor show lower baseline firing and focal nicotine-induced increases in firing.

**Fig 4.**
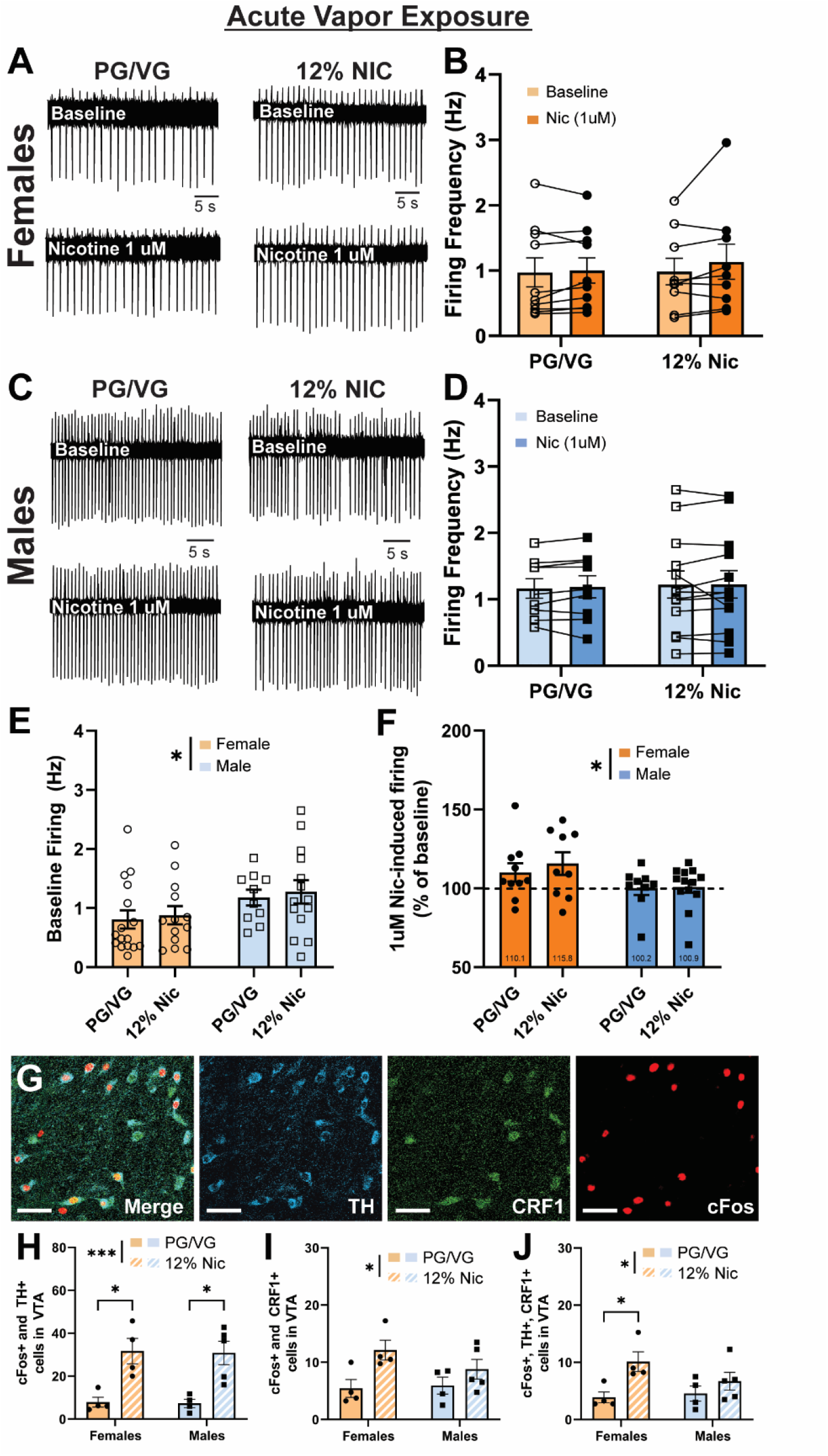
Spontaneous firing in ^VTA-NAc^CRF1 neurons and population activity in VTA following acute vapor exposure. **(A)** Spontaneous firing in ^VTA-NAc^CRF1 neurons from female mice exposed to acute PG/VG (left) or 12% nicotine (right) at baseline (top) and after 1 μM nicotine application (bottom). **(B)** Spontaneous firing frequency in ^VTA-NAc^CRF1 neurons from PG/VG (N=10 cells, N=5 mice) or 12% nicotine (N=9 cells, N=5 mice) exposed female mice at baseline (open circle) and after 1 μM nicotine application (filled circles). **(C)** Spontaneous firing in ^VTA-NAc^CRF1 neurons from male mice exposed to PG/VG (left) or 12% nicotine (right) at baseline and after 1 μM nicotine application. **(D)** Spontaneous firing frequency in ^VTA-NAc^CRF1 neurons from PG/VG (N=9 cells, N=4 mice) or 12% nicotine (N=13 cells, N=4 mice) exposed male mice at baseline (open squares) and after 1 μM nicotine application (filled squares). **(E)** Baseline firing in ^VTA-NAc^CRF1 neurons from females (circles) and males (squares) exposed to PG/VG (Female N=16 cells, N=5 mice; Male N=10 cells, N=4 mice) or 12% nicotine (Female N=13 cells, N=5 mice; Male N=14 cells, N=4 mice). There was a main effect of sex (*p=0.0282, F (1, 49) = 5.114) by 2-way ANOVA. **(F)** 1 μM nicotine-induced change in firing in ^VTA-NAc^CRF1 neurons from female and male mice exposed to PG/VG (Female N=10 cells, N=5 mice; Male N=9 cells, N=4 mice) or 12% nicotine (Female N=9 cells, N=5 mice; Male N=13 cells, N=4 mice). There was a main effect of sex (*p=0.0253, F (1, 37) = 5.437) by 2-way ANOVA. **(G)** Representative image of merged, Th expression, CRF1 expression, and cFos expression in the VTA, scale bar = 50 μm. **(H)** Number of VTA neurons that express cFos and Th in females and males exposed to acute PG/VG (Female N=4, Male N=4) or acute 12% nicotine (Female N=4, Male N=5). 12% nicotine exposed animals showed greater expression (main effect of vapor content ***p=0.0002, F (1, 13) = 27.47, Female: PG/VG vs Nic *p=0.0140, Male: PG/VG vs Nic *p=0.0107, 2-way ANOVA post hoc Tukey’s). **(I)** Number of VTA neurons that express cFos and CRF1 in females and males exposed to acute PG/VG (Female N=4, Male N=4) or acute 12% nicotine (Female N=4, Male N=5). 12% nicotine exposed animals showed greater expression (main effect of vapor content *p=0.0127, F (1, 13) = 8.350, 2-way ANOVA). **(J)** Number of VTA neurons that express cFos, Th, CRF1 in females and males exposed to acute PG/VG (Female N=4, Male N=4) or acute 12% nicotine (Female N=4, Male N=5). 12% nicotine exposed animals showed greater expression (main effect of vapor content *p=0.0129, F (1, 13) = 8.283, Female: PG/VG vs Nic *p=0.0482, 2-way ANOVA post hoc Tukey’s). Data are represented as mean ± SEM. See also Figure S2.

### The effects of acute vapor exposure on VTA subpopulation activity

Acute 12% nicotine vapor exposure (**Fig 4G**) did not alter the number of TH+ or CRF1+ neurons in the VTA of either sex (**Supp Fig 2A-B**), however, the number of cFos+ neurons was increased (main effect of vapor content *p=0.0217, F (1, 13) = 6.796, 2-way ANOVA; **Supp Fig 2C**). cFos expression was increased in the TH+ (dopaminergic) VTA population in both sexes exposed to 12% nicotine compared to PG/VG (main effect of vapor content ***p=0.0002, F (1, 13) = 27.47, Female: PG/VG vs Nic *p=0.0140, Male: PG/VG vs Nic *p=0.0107, 2-way ANOVA post hoc Tukey’s; **Fig 4H**). cFos expression in the CRF1+ VTA population was significantly higher in acute 12% nicotine-exposed mice as compared to PG/VG-exposed mice, however, post hoc testing did not reach significance (main effect of vapor content *p=0.0127, F (1, 13) = 8.350, 2-way ANOVA; **Fig 4I**). cFos expression in TH+ and CRF1+ co-expressing neurons was also increased in 12% nicotine vapor-exposed animals, but post hoc comparisons were only significant for females (main effect of vapor content *p=0.0129, F (1, 13) = 8.283, Female: PG/VG vs Nic *p=0.0482, 2-way ANOVA post hoc Tukey’s; **Fig 4J**). Expression of cFos in each of these neuronal populations was not statistically correlated with serum nicotine levels in females, males, or both sexes combined (**Supp Fig 2D-F**). Overall, these data indicate that acute nicotine vapor exposure increases activity in VTA dopamine and VTA CRF1 neuronal populations of both sexes but with more profound effects in females.

### The effect of chronic vapor exposure on inhibitory signaling of ^VTA-NAc^CRF1 neurons

We exposed CRF1-GFP mice to chronic vapor (**Fig 5A, left**) and found no significant differences in membrane properties in ^VTA-NAc^CRF1 neurons (**Fig 5A, right**). In females (**Fig 5B**), focal nicotine (1 μM) did not change sIPSC frequency (**Fig 5C, left and Fig 5G, left**) but increased sIPSC amplitude, mainly driven by the PG/VG group (main effect of focal nicotine *p=0.0147, F (1, 18) = 7.273, PV/VG: baseline vs 1uM Nic *p=0.0155, RM 2-way ANOVA post hoc Sidak’s; **Fig 5C, right**). However, when the 1 μM nicotine-induced change in sIPSC amplitude was normalized to baseline, we did not observe a significant increase in either sex (**Fig 5G, right**). In males (**Fig 5D**), focal nicotine (1 μM) produced no change in sIPSC frequency (**Fig 5E, left and Fig 5G, left**) but reduced sIPSC frequency in 12% nicotine-exposed as compared to PG/VG-exposed mice (main effect of vapor content *p=0.0169, F (1, 26) = 6.521, RM 2-way ANOVA; **Fig 5E, left**). Focal nicotine increased sIPSC amplitude but post hoc Sidak’s test did not show significance between any specific groups (main effect of focal nicotine *p=0.0467, F (1, 26) = 4.361, RM 2-way ANOVA; **Fig 5E, right**). When the 1μM nicotine-induced change in sIPSC frequency and amplitude were normalized to baseline there was no significant difference between sexes from either PG/VG or 12% nicotine groups (**Fig 5G**). Baseline sIPSC frequency was lower in 12% nicotine-exposed mice, likely driven by males, however post hoc Tukey’s did not yield significant differences between groups (main effect of vapor content *p=0.0385, F (1, 52) = 4.508, 2-way ANOVA; **Fig 5F, left**). Baseline sIPSC amplitude was not altered in either sex exposed to PG/VG or 12% nicotine (**Fig 5F, right**). Focal nicotine induced a tonic inhibitory current in ^VTA-NAc^CRF1 neurons of both sexes exposed to either PG/VG or 12% nicotine (**Fig 5H-I**) that was partially reversed with 100 μM gabazine (**Fig 5J**). These data suggest that chronic nicotine vapor exposure reduced presynaptic phasic inhibition in ^VTA-NAc^CRF1 neurons from males, increased postsynaptic phasic inhibition in both sexes, and the sex difference in tonic inhibition observed following acute nicotine vapor exposure was no longer present.

**Fig 5.**
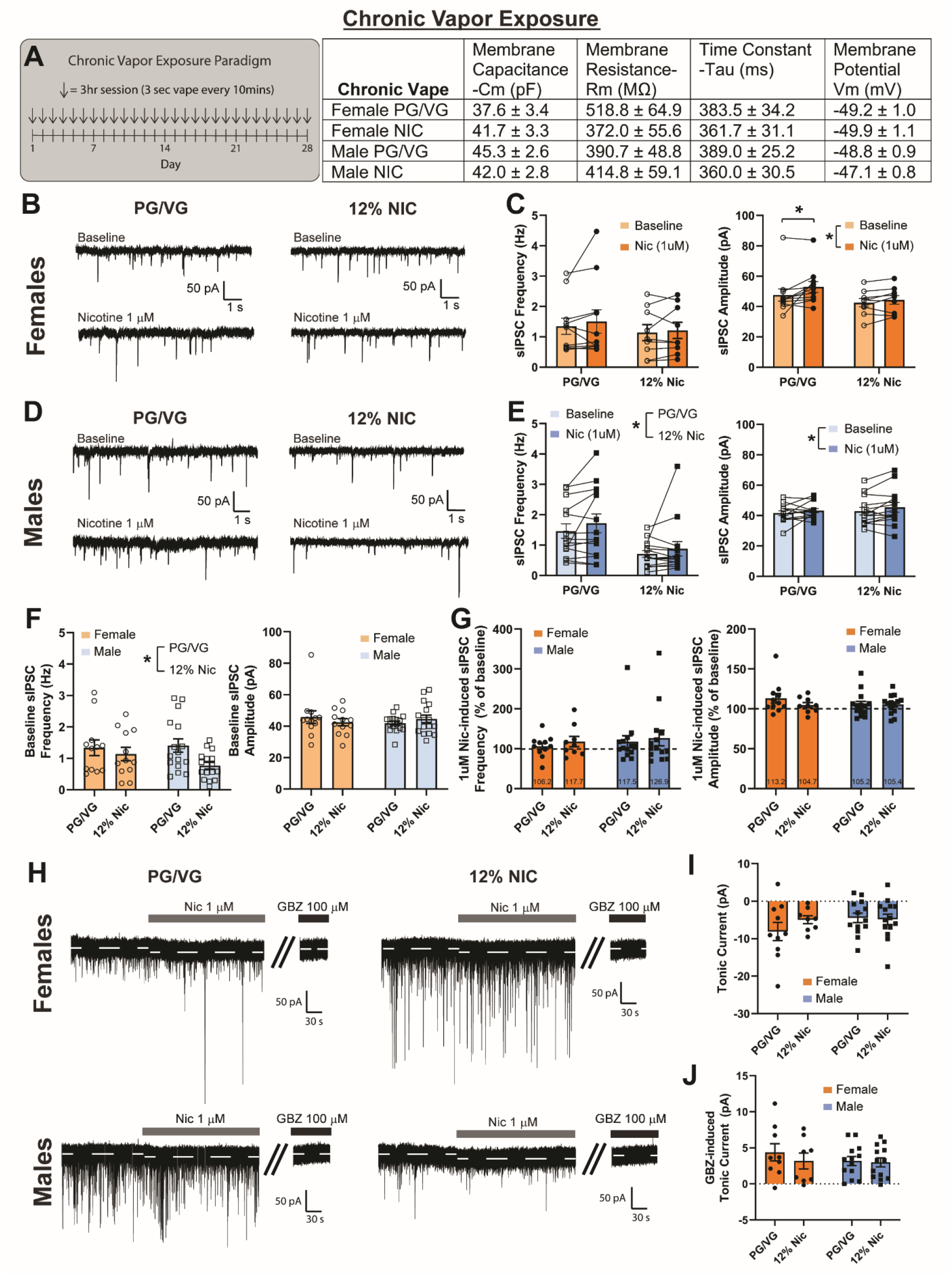
Inhibitory signaling in ^VTA-NAc^CRF1 neurons following chronic vapor exposure. **(A)** Chronic vapor exposure paradigm (left) and membrane properties (right) of ^VTA-NAc^CRF1 neurons from male and female mice exposed to chronic PG/VG (Female N=24 cells, N=4 mice; Male N= 33 cells, N=6 mice) or 12% nicotine vapor (Female N=25 cells, N=5 mice; Male N= 32 cells, N=6 mice). **(B)** Spontaneous inhibitory post-synaptic currents (sIPSCs) in ^VTA-NAc^CRF1 neurons from female mice exposed to PG/VG (left) or 12% nicotine (right) at baseline and after 1 μM nicotine application. **(C)** sIPSC frequency (left) and amplitude (right) in ^VTA-NAc^CRF1 neurons from PG/VG (N=11 cells, N=4 mice) or 12% nicotine (N=9 cells, N=5 mice) exposed female mice at baseline and after 1 μM nicotine application. Focal nicotine increased sIPSC amplitude (main effect of focal nicotine *p=0.0147, F (1, 18) = 7.273, RM 2-way ANOVA), mainly driven by the PG/VG group (PV/VG: baseline vs 1uM Nic *p=0.0155, post hoc Sidak’s). **(D)** Spontaneous inhibitory post-synaptic currents (sIPSCs) in ^VTA-NAc^CRF1 neurons from male mice exposed to PG/VG (left) or 12% nicotine (right) at baseline and after 1 μM nicotine application. **(E)** sIPSC frequency (left) and amplitude (right) in ^VTA-NAc^CRF1 neurons from PG/VG (N=14 cells, N=6 mice) or 12% nicotine (N=14 cells, N=6 mice) exposed male mice at baseline and after 1 μM nicotine application. sIPSC frequency reduced in 12% nicotine-exposed animals (main effect of vapor content *p=0.0169, F (1, 26) = 6.521, RM 2-way ANOVA) and focal nicotine increased sIPSC amplitude (main effect of focal nicotine *p=0.0467, F (1, 26) = 4.361, RM 2-way ANOVA). **(F)** Baseline sIPSC frequency (left) and amplitude (right) from female and male mice exposed to either PG/VG (Female N=13 cells, N=4 mice; Male N=16 cells, N=6 mice) or 12% nicotine (Female N=12 cells, N=5 mice; Male N=16 cells, N=6 mice). 12% nicotine exposed animals show lower baseline sIPSC frequency (main effect of vapor content *p=0.0385, F (1, 52) = 4.508, 2-way ANOVA). **(G)** 1 μM nicotine-induced change in sIPSC frequency (left) and amplitude (right) normalized to baseline in ^VTA-NAc^CRF1 neurons from female and male mice exposed to PG/VG (Female N=11 cells, N=4 mice; Male N=14 cells, N=6 mice) or 12% nicotine (Female N=9 cells, N=5 mice; Male N=14 cells, N=6 mice). **(H)** Example traces of 1 μM nicotine-induced and gabazine (GBZ) reversed tonic current in ^VTA-NAc^CRF1 neurons from female (top) and male (bottom) mice exposed to PG/VG (left) or 12% nicotine (right). **(I)** 1 μM nicotine-induced tonic current in ^VTA-NAc^CRF1 neurons from female and male mice exposed to PG/VG (Female N=10 cells, N=3 mice; Male N=13 cells, N=6 mice) or 12% nicotine (Female N=8 cells, N=5 mice; Male N=14 cells, N=6 mice). **(J)** Gabazine (GBZ, 100 μM) reversal of tonic current in ^VTA-NAc^CRF1 neurons from female and male mice exposed to PG/VG (Female N=9 cells, N=3 mice; Male N=12 cells, N=6 mice) or 12% nicotine (Female N=8 cells, N=5 mice; Male N=13 cells, N=5 mice). Data are represented as mean ± SEM.

### The effect of chronic vapor exposure on spontaneous firing of ^VTA-NAc^CRF1 neurons

We performed cell-attached recordings in female (**Fig 6A, B**) and male (**Fig 6C, D**) mice exposed to chronic PG/VG or 12% nicotine vapor and observed no differences in baseline firing in either vapor condition (**Fig 6E**). Focal nicotine (1 μM) did not significantly change spontaneous firing (vapor content x sex interaction *p=0.0452, F (1, 38) = 4.290, 2-way ANOVA; **Fig 6F**) and the normalized percent change of firing was not significantly altered in females or males in either vapor group. These data suggest that sex differences in baseline firing and effects of focal nicotine on firing in ^VTA-NAc^CRF1 neurons following acute exposure is reversed following chronic exposure to PG/VG or 12% nicotine vapor.

**Fig 6.**
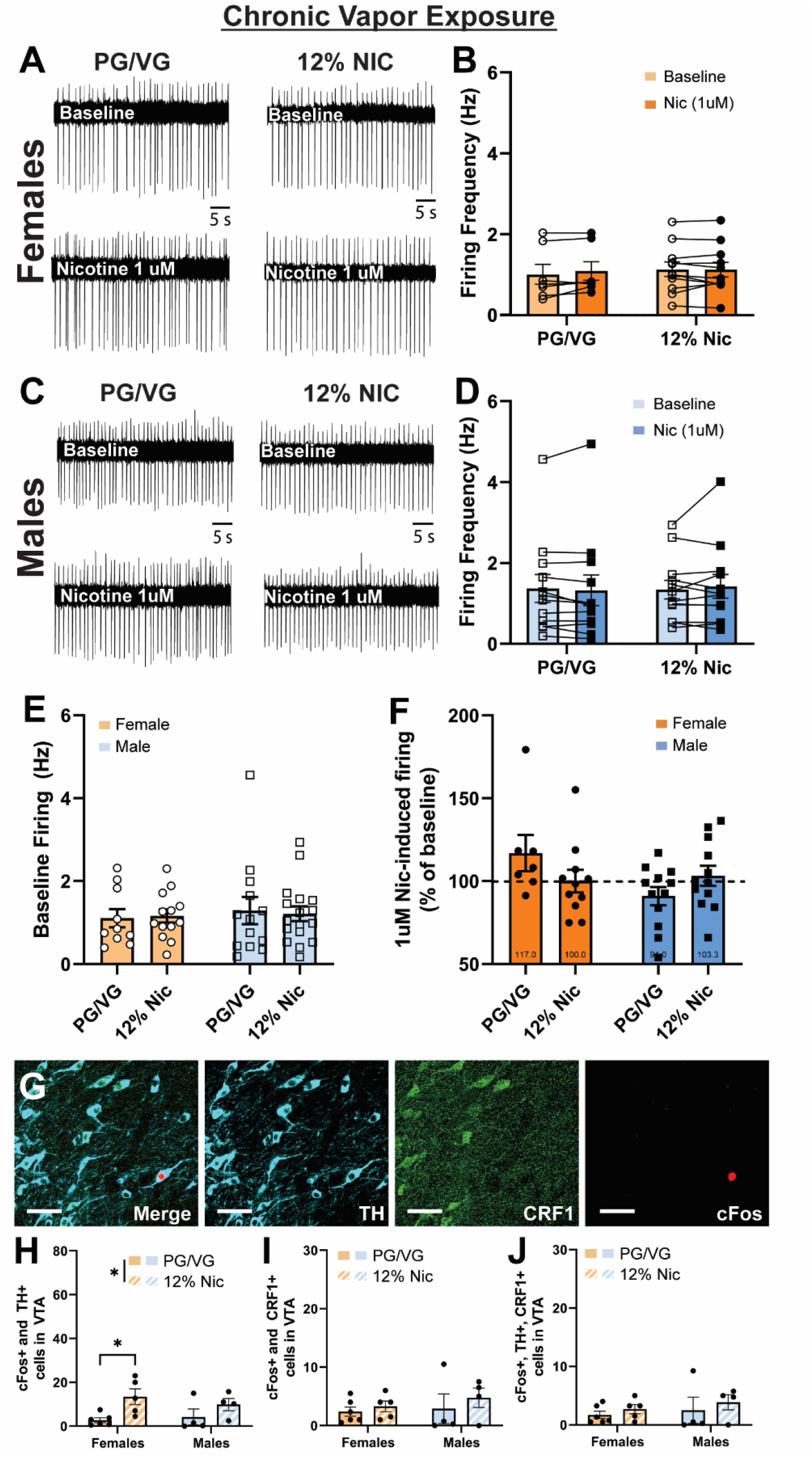
Spontaneous firing in ^VTA-NAc^CRF1 neurons and population activity in VTA following chronic vapor exposure. **(A)** Spontaneous firing in ^VTA-NAc^CRF1 neurons from female mice exposed to chronic PG/VG (left) or 12% nicotine (right) at baseline and after 1 μM nicotine application. **(B)** Spontaneous firing frequency in ^VTA-NAc^CRF1 neurons from chronic PG/VG (N=7 cells, N=4 mice) or 12% nicotine (N=11 cells, N=5 mice) exposed female mice at baseline (open circles) and after 1 μM nicotine application (filled circles). **(C)** Spontaneous firing in ^VTA-NAc^CRF1 neurons from male mice exposed to chronic PG/VG (left) or 12% nicotine (right) at baseline and after 1 μM nicotine application. **(D)** Spontaneous firing frequency in ^VTA-NAc^CRF1 neurons from chronic PG/VG (N=12 cells, N=5 mice) 12% nicotine (N=12 cells, N=4 mice) exposed male mice at baseline (open squares) and after 1 μM nicotine application (filled squares). **(E)** Baseline firing in ^VTA-NAc^CRF1 neurons from females (circles) and males (squares) exposed to chronic PG/VG (Female N=10 cells, N=4 mice; Male N=13 cells, N=5 mice) or 12% nicotine (Female N=13 cells, N=5 mice; Male N=17 cells, N=5 mice). **(F)** 1 μM nicotine-induced change in firing in ^VTA-NAc^CRF1 neurons from female and male mice exposed to chronic PG/VG (Female N=7 cells, N=4 mice; Male N=12 cells, N=5 mice) or 12% nicotine (Female N=11 cells, N=5 mice; Male N=12 cells, N=4 mice). **(G)** Representative image of merged, Th expression, CRF1 expression, and cFos expression in the VTA. Scale bar = 50 μm. **(H)** Number of VTA neurons that express cFos and Th in females and males exposed to chronic PG/VG (Female N=6, Male N=4) or acute 12% nicotine (Female N=5, Male N=4). 12% nicotine exposed animals showed greater expression (main effect of vapor content *p=0.0102, F (1, 15) = 8.615, female: PG/VG vs nicotine *p=0.0443, 2-way ANOVA post hoc Tukey’s). **(I)** Number of VTA neurons that express cFos and CRF1 in females and males exposed to chronic PG/VG (Female N=6, Male N=4) or acute 12% nicotine (Female N=5, Male N=4). **(J)** Number of VTA neurons that express cFos, Th, CRF1 in females and males exposed to chronic PG/VG (Female N=6, Male N=4) or acute 12% nicotine (Female N=5, Male N=4). Data are represented as mean ± SEM. See also Figure S3

### The effect of chronic vapor exposure on VTA subpopulation activity

Chronic 12% nicotine vapor exposure (**Fig 6G**) did not alter the number of TH+ or CRF1+ neurons in the VTA of either sex (**Supp Fig 3A-B**) but did increase the number of cFos+ neurons (main effect of vapor content *p=0.0446, F (1, 15) = 4.804, 2-way ANOVA; **Supp Fig 3C**). cFos expression in the TH+ VTA population was increased in 12% nicotine-exposed mice compared to PG/VG-exposed mice and post hoc tests showed a significant difference between vapor groups in females but not males (main effect of vapor content *p=0.0102, F (1, 15) = 8.615; Female: PG/VG vs nicotine *p=0.0443, 2-way ANOVA post hoc Tukey’s; **Fig 6H**). There was no difference in cFos expression in the CRF1+ or the TH+/CRF1+ VTA population in either sex from both vapor groups (**Fig 6I-J**). Expression of cFos in each of these neuronal populations was positively correlated with serum nicotine levels in females, males, and both sexes combined, however, the correlation were not statistically significant (**Supp Fig 3D-F**). These data indicate that chronic nicotine vapor exposure maintains elevated activity in VTA dopaminergic neurons, especially in females. However, the increased activity in CRF1+ neurons following acute exposure is reduced to levels similar to PG/VG after chronic exposure.

## Discussion

CRF1 neurons in the VTA are predominately dopaminergic and project to the NAc, suggesting involvement in the mesolimbic reward pathway. ^VTA-NAc^CRF1 neurons showed greater phasic inhibition in naïve females, greater focal nicotine-induced firing in naïve males, and both sexes displayed focal nicotine-induced tonic inhibition. Following acute nicotine vapor exposure, phasic inhibition was largely unaffected, but focal nicotine-induced tonic inhibition was enhanced in females and reduced in males. Acute nicotine exposure did not affect firing in ^VTA-NAc^CRF1 neurons, but females showed lower baseline firing and increased focal nicotine-induced firing. Following acute nicotine vapor exposure, activity of the CRF1 dopaminergic VTA population increased in both sexes, but especially in females. In contrast, chronic nicotine vapor exposure reduced basal phasic inhibition in ^VTA-NAc^CRF1 neurons in both sexes and the sex-specific bidirectional change in tonic inhibition was no longer observed. Additionally, activity of the CRF1 dopaminergic VTA population was no longer elevated following chronic nicotine vapor exposure. Collectively, these findings demonstrate sex-specific differences in inhibitory control and nicotine sensitivity in ^VTA-NAc^CRF1 neurons from naïve mice, and sex-specific changes in activity and inhibitory control following acute nicotine vapor exposure that were lost following chronic exposure. Taken together, these findings shed light on important sex- and exposure-dependent changes in the mesolimbic CRF1 population and how electronic nicotine vapor selectively impacts reward and stress circuitry in males and females.

ENDS use has increased dramatically, however, the effects of nicotine vapor on brain reward and stress circuitry remain unclear. Previous studies have used alternative forms of nicotine delivery like subcutaneous minipumps^24, 36^, drinking water^37, 38^, intraperitoneal^39–41^, or intravenous^4, 5^ injections. This study used a nicotine vapor inhalation model that allows nicotine to reach the brain on a timescale similar to nicotine from cigarettes^42^, which is a primary determinant of nicotine reinforcement^43^. Additionally, vapor liquids contain higher nicotine concentrations, which may cause differential engagement of the reward circuitry. Our previous study found that a single 3-second 12% nicotine vapor produced mouse serum nicotine levels^33^ that are comparable to human serum levels in cigarette smokers^44^ and electronic cigarette users^45^. Additionally, serum cotinine levels in mice following acute nicotine vapor exposure (Supp. Fig 1A right) were comparable to those found in humans who smoke cigarettes daily or use e-cigarettes daily^46^. However, with repeated nicotine vapor administrations (acute and chronic exposure paradigms), we did observe serum nicotine levels higher than what have been reported in humans. This discrepancy can be due to a variety of potential factors including the species’ different nicotine metabolism rates (mice have faster nicotine metabolism than humans), route of inhalation and absorption (mice inhale the vapor mainly through the nose whereas humans inhale vapor through the mouth), time of sample collection (mouse samples in our study collected immediately following last vape session vs human sample collection can vary depending on the study), and naturalistic mouse behavior (potential ingestion of nicotine deposits on fur when grooming). Although nicotine vapor delivery models human nicotine vapor inhalation, human drug use involves a component of volition. This study uses a passive form of vapor exposure but future experiments incorporating self-administration will model animals’ motivation to seek nicotine.

Clinical reports have found that women, more than men, report stress as a major factor promoting nicotine use^14^. In addition, sex differences in nicotine metabolism^47, 48^ could contribute to neurobiological effects. These human sex differences highlight the gap in knowledge on nicotine effects in females and the need for female subjects in pre-clinical research. We examined ^VTA-NAc^CRF1 neurons in both sexes and found that females show higher basal phasic inhibition and enhanced tonic inhibition after exposure to acute nicotine vapor compared to male counterparts. These data indicate that female VTA CRF1 neurons are under greater inhibitory control which may explain the female-specific decrease in dopamine release in the nucleus accumbens with administration of the nicotinic receptor antagonist, mecamylamine^49^. Overexpression of CRF in the nucleus accumbens has also been shown to enhance the reinforcing effects of nicotine in females^25^, thus future experiments examining the effects of CRF in both female and male ^VTA-NAc^CRF1 neurons may shed light on how stress can affect reward processing in the VTA.

Following acute nicotine vapor exposure, VTA CRF1 neuronal activity (cFos) was increased in both sexes and focal nicotine increased firing in males, consistent with previous studies in VTA dopamine neurons^4, 50–52^. However, few studies have examined focal nicotine effects in females. Nicotine activates nicotinic acetylcholine receptors (nAChRs) on dopamine neurons to drive increases in firing and dopamine release in the NAc^53^. VTA DA neurons are also modulated by nicotine’s activation of α7 nAChRs on glutamatergic inputs and activation of α4β2 nAChRs on GABAergic inputs. Phasic GABAergic inhibitory inputs are transiently increased but quickly desensitized leading to an overall increase in excitability in the postsynaptic VTA dopamine neuron^1, 2^. Additionally, there’s evidence that tonic inhibition is also present in VTA neurons^54^ however, tonic inhibition in the context of nicotine is understudied. Although we demonstrate that VTA CRF1 neurons are primarily dopaminergic, phasic inhibitory signaling in ^VTA-NAc^CRF1 neurons was not sensitive to either focal or *in vivo* nicotine, in contrast to previous work in VTA dopamine neurons. This difference may be due to differences in species (rat vs mouse), sex (male vs both sexes), and/or age (adolescent vs adult). However, our data suggest that modulation of the ^VTA-NAc^CRF1 neuron population in response to nicotine exposure is different than what is reported in the VTA dopaminergic population. These differences could underly potentially distinct roles in stress/anxiety and/or addiction.

VTA dopamine neurons are modulated by both phasic^1, 24^ and tonic^54^ inhibition. Phasic inhibition in ^VTA-NAc^CRF1 neurons was not significantly altered by acute or chronic electronic nicotine vapor exposure or focal nicotine application. In contrast, focal nicotine induced a tonic inhibition in both naïve female and male ^VTA-NAc^CRF1 neurons and following acute nicotine vapor exposure, tonic inhibition was enhanced in females and reduced in males. The enhanced tonic inhibition in females can dampen glutamatergic inputs and thus, decrease reward signaling whereas the reduced tonic inhibition in males could enhance glutamatergic inputs and drive reward signaling. These findings may be the mechanistic basis underlying sex difference in the motivation for nicotine seeking where women are more likely to seek nicotine to relieve stress and men are more likely to seek nicotine for its rewarding properties. Additionally, these findings suggest that nicotine’s effect on excitability of the ^VTA-NAc^CRF1 population may be modulated more by tonic inhibition than phasic inhibition as previously observed in VTA dopamine neurons^1, 24^. Interestingly, the bidirectional sex difference in focal nicotine-induced tonic inhibition was no longer observed following chronic nicotine vapor, suggesting the occurrence of neuroplastic adaptations that dampen the ability of tonic inhibition to regulate ^VTA-NAc^CRF1 neuronal activity with chronic exposure. Distinct changes following acute but not repeated nicotine vapor exposure are consistent with previous work in the central amygdala^33^. The differential effects of nicotine on tonic inhibition, and thus, the excitability of ^VTA-NAc^CRF1 neurons may play a role in the modulation of the reward circuit and the development and maintenance of addiction.

Neuronal activity in this study was measured by electrophysiological recordings of spontaneous firing to assess dynamic changes in individual neuron activity and immunohistochemical expression of the activity marker, cFos, to assess activity across neuronal populations. Acute nicotine vapor exposure did not alter spontaneous firing but increased cFos expression. These results may at first seem contradictory but rather underline the importance of understanding the temporal aspect of nicotine’s effects. Mice used for both methods of measuring neuronal activity were sacrificed immediately following the last vapor session. However, spontaneous firing was measured following slice preparation and at least an hour-long incubation in artificial cerebral spinal fluid (aCSF) when neurons are potentially in cellular withdrawal with variable levels of nicotine remaining in the brain. Whereas cFos expression reflects a snapshot of neuronal activity ~30-60 minutes prior to the time of sacrifice^55^ while the animals are actively being exposed to the vapor. The cFos increase following acute and decrease following chronic nicotine vapor exposure is consistent with previous studies examining the patterns of activity following systemic nicotine exposure^51^.

The CRF/CRF1 system is involved in reward processing as previous studies report CRF-induced increases in dopamine firing^27, 28^, coordination of reward behavior^29^, and reinforcement of drugs of abuse like cocaine^17^. In the context of nicotine, studies have shown increased CRF mRNA in the VTA following chronic nicotine exposure ^24^ and blocking CRF1 reduced reinstatement of nicotine seeking^31^. Activation of the cell bodies of VTA dopamine CRF1 neurons have been shown to coordinate reward reinforcement behavior and activation of their terminals in the NAc core increase dopamine release^29^. Additionally, CRF peptide increases VTA dopamine neuron firing via CRF1^27, 28^ and knockout of CRF1 in midbrain dopamine neurons have been shown to increase anxiety-like behavior^30^. Previous studies have found *Crf1* mRNA expression in GABAergic neurons in the VTA^28^, however the location of the protein expression (dendrites/cell body vs terminals) and their role in stress/anxiety and nicotine addiction is still unclear. We targeted CRF1 neurons in the VTA to better understand how this population is involved in nicotine reward processing. We found that these neurons are mainly dopaminergic and project to the nucleus accumbens, consistent with previous reports^29^. However, ^VTA-NAc^CRF1 neurons do not respond to nicotine in the same way as canonical VTA dopamine neurons and given the relevance of CRF1 in stress signaling, these neurons may be uniquely modulated in the context of stress and anxiety. Future studies examining how nicotine reward processing is affected by stress and anxiety and how anxiety-like behaviors are affected by nicotine exposure are needed to better understand the intersection of divergent (or convergent) processing in CRF1 VTA neurons in different behavioral conditions.

Collectively, these studies provide important information on how a neuronal population relevant to stress is involved in encoding nicotine reward and how *in vivo* exposure to acute and chronic electronic nicotine vapor exposure produces different effects on activity and inhibitory control of the VTA CRF1 population in females and males. These sex- and exposure-dependent changes in the mesolimbic CRF1 population add to our understanding of the neurobiological underpinnings involved in the development of nicotine addiction. Our electronic nicotine delivery approach provides model of nicotine consumption in human vape users. It’s imperative that we continue to study how nicotine vapor affects the brain and engages specific reward pathways to better understand nicotine addiction and aid in the development of therapeutics to prevent and mitigate nicotine addiction.

## Materials and Methods

### Animals

Adult CRF1-GFP^32^ or CRF1-Cre mice on a C57/BL6J background were group-housed in a temperature- and humidity-controlled 12hr light/dark (7am lights on) facility with ad libitum food and water access. All experimental procedures were approved by the UNC Institutional Animal Care and Use Committee.

### Immunohistochemistry

Mice were perfused and tissue sectioned and processed as previously described^33^. Sections were incubated with primary antibodies: mouse anti-TH (1:1000; T1229, Sigma), chicken anti-GFP (1:1000; ab13970, Abcam), rabbit anti-cFos (1:3000; ABE457, Millipore) and secondary antibodies: Alexa 555 goat anti-mouse (1:250; A21424, Invitrogen), Alexa 647 goat anti-mouse (1:200; 115-605-003, JacksonImmuno), Alexa 488 donkey anti-chicken (1:700; 703-545-155, JacksonImmuno) or goat anti-rabbit horse radish peroxidase (1:200; ab6721, Abcam) followed by tyramide-conjugated Cy3 (1:50) diluted in TSA amplification diluents (Akoya Biosciences, NEL741001KT). Sections were mounted, cover-slipped, and imaged on a confocal microscope (Leica SP8). Quantification was performed by experimenters blinded to experimental groups using ImageJ (NIH) and counts from two sections/animal were averaged.

### Stereotaxic Intracranial Microinjection

Mice were anesthetized with isoflurane (4% induction, 2-3% maintenance) for stereotaxic (Kopf Instruments) intracranial injections using microsyringe (Hamilton, 88411) syringe pump infusion of 100 nl/min into the target region. For anatomy, CRF1-cre mice were injected with 500 nL/hemisphere of AAV5-hSyn-DIO-eGFP (50457-AAV5; titer ≥ 7×10¹² vg/mL, Addgene) into the VTA (ML ±0.60, AP −3.2, DV −4.5). For electrophysiology, CRF1-GFP mice were injected with 250 nL/hemisphere red retrograde beads (Lumafluor) into the nucleus accumbens (ML ±0.65, AP +1.48, DV −4.75).

### Slice Electrophysiology

Immediately after completion of last vapor exposure, mice were rapidly decapitated and brains extracted into sucrose solution containing (in mM): sucrose 206.0; KCl 2.5; CaCl_2_ 0.5; MgCl_2_ 7.0; NaH_2_PO_4_ 1.2; NaHCO_3_ 26; glucose 5.0; HEPES 5. Slices (250-300 μM) were incubated in oxygenated (95% O_2_/5% CO_2_) artificial cerebrospinal fluid (aCSF) containing (in mM): NaCl 130; KCL 3.5; CaCl_2_ 2; MgSO_4_7H_2_O 1.5; NaH_2_PO_4_H_2_O 1.25; NaHCO_3_ 24; Glucose 10 at 37°C (30 min) and room temperature (30 min). Recordings were performed 1-8 hours following decapitation with pipettes (4-7 MΩ) filled with internal solution (in mM) KCl 145; EGTA 5; MgCl_2_ 5; HEPES 10; Na-ATP 2; Na-GTP 0.2. Inhibitory transmission was measured using whole-cell voltage-clamp (V_hold_= −60 mV) recordings with receptor antagonists (20 μM DNQX, 50 μM AP5, 1 μM CGP 52432) in the superfused aCSF. Recordings were obtained with a Multiclamp 700B amplifier (Molecular Devices), digitized (Digidata 1440A; Molecular Devices), and stored on a computer (pClamp 10 software; Molecular Devices). Nicotine (1 μM) and gabazine (100 μM; SR 955531) were applied by a y-tube positioned in close proximity to the cell on the surface of the slice. Nicotine applied by y-tube is referred to as ‘focal nicotine’. Firing frequency was quantified by Clampfit 11.1 (Molecular Devices). Spontaneous inhibitory postsynaptic currents (sIPSCs), or phasic inhibition, were analyzed using MiniAnalysis (Synaptosoft). Baseline sIPSCs were analyzed after >3 minutes of superfused receptor antagonists. Focal nicotine (1 μM)-induced sIPSCs were analyzed for ~2 minutes as previously reported ^1^. Baseline and experimental sIPSCs were analyzed from stable recording periods with ≥60 events. The mean holding current was measured using a Gaussian fit to an all-points histogram of the holding current over a 5 second period as previously described^34^. Tonic current, or tonic inhibition, was calculated by taking the difference in mean holding current before and after y-tube drug application.

All electrophysiological measures are reported as values from individual cells and recordings from multiple cells are taken from each animal. Data are presented as raw values or normalized data (percent change) to account for variability in baseline. Percent change from baseline is calculated by (drug/baseline)*100, thus values above 100% are increases from baseline and values below 100% are decreases from baseline.

### Electronic Nicotine Vapor Exposure

Mice were placed in airtight vacuum-controlled chambers (~1 L/min air circulation). Vaporizers (95 watt, SVS200, Scientific Vapor) heat up solutions in e-vape tanks (Baby Beast Brother, Smok) containing 12% (v/v) (−)-nicotine free base in a 50/50 (v/v) propylene glycol/vegetable glycerol (PG/VG) or PG/VG solution ^33^. The 12% (120 mg/ml) nicotine concentration reflects a 2-fold increase from commercial e-liquids (i.e. JUUL 59 mg/ml), accounting for the passive inhalation from chamber and higher metabolism of nicotine in mice. E-vape controllers (SSV-1, La Jolla Alcohol Research) triggered pre-set vapor delivery. Acute vapor exposure was composed of 3-sec long vapor deliveries every 10 minutes over the course of a 3 hour session. Chronic vapor exposure is composed of 28 consecutive days of daily 3-hour sessions of 3 sec vape every 10 minutes. In both acute and repeated exposure paradigms female and male mice were exposed to either PG/VG control or 12% nicotine in PG/VG.

### Nicotine and Cotinine Serum Analysis

Trunk blood was collected immediately following final vapor exposure. Samples were centrifuged and serum was collected and stored at −20°C. Samples were analyzed for metabolites using liquid chromatography-tandem mass spectrometry (LC-MS/MS) as previously described^35^.

### Statistical Analysis

Statistical analysis and data visualization were conducted using Prism 9.0 (GraphPad). Data were analyzed and compared using unpaired two-tailed t-tests (with Welch’s correction where appropriate), one-sample t-tests (theoretical mean = 100), and two-way ANOVAs with post-hoc Sidak’s or Tukey’s and repeated measures when there is a significant interaction or main effect. Variance was assessed using F test or Bartlett’s test and normality was assessed using Kolmogorov-Smirnov test. All data are expressed as mean ± SEM with p<0.05 as the criterion for statistical significance. For detailed statistical reporting see Supplemental Table.

## Acknowledgements

The authors would like to thank Dr. Charles R. Esther and Tara Nicole Guhr Lee for their help with the serum nicotine and cotinine analysis. We would also like to thank Maria Echeveste Sanchez for technical assistance. This work was funded by F31-DA-053064 (MZ), T32-NS-007431 (MZ), AA-026858 (MAH), and AA-011605 (MAH).

## Author contributions

MZ and MAH conceptualized the project and designed the experiments. MZ performed surgeries and performed electrophysiology experiments and analysis. MZ, NGR, and JVJ performed behavior experiments and IHC experiments. MZ drafted manuscript. MZ and MAH edited manuscript. All authors have read and approved the final manuscript.

---

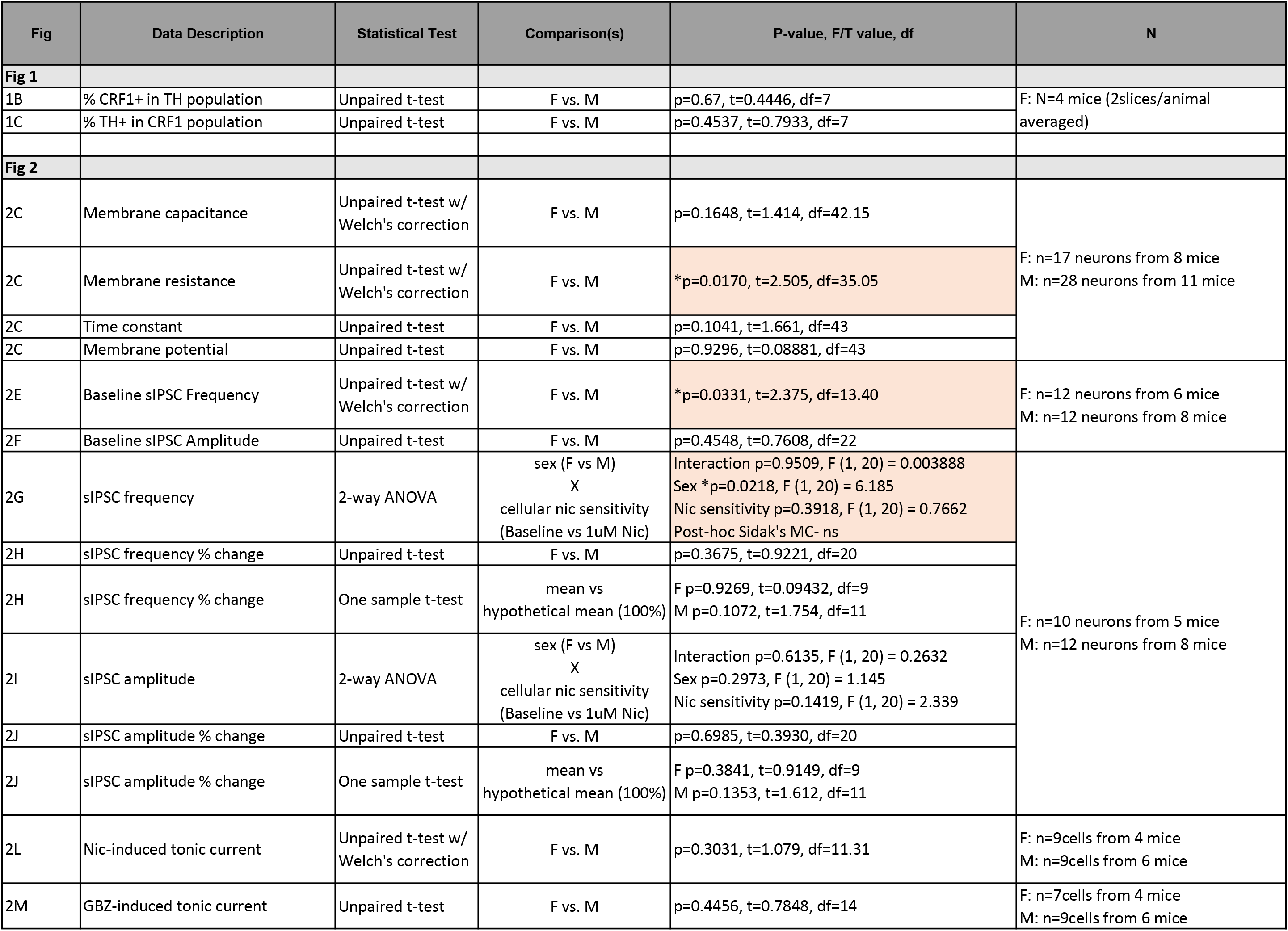
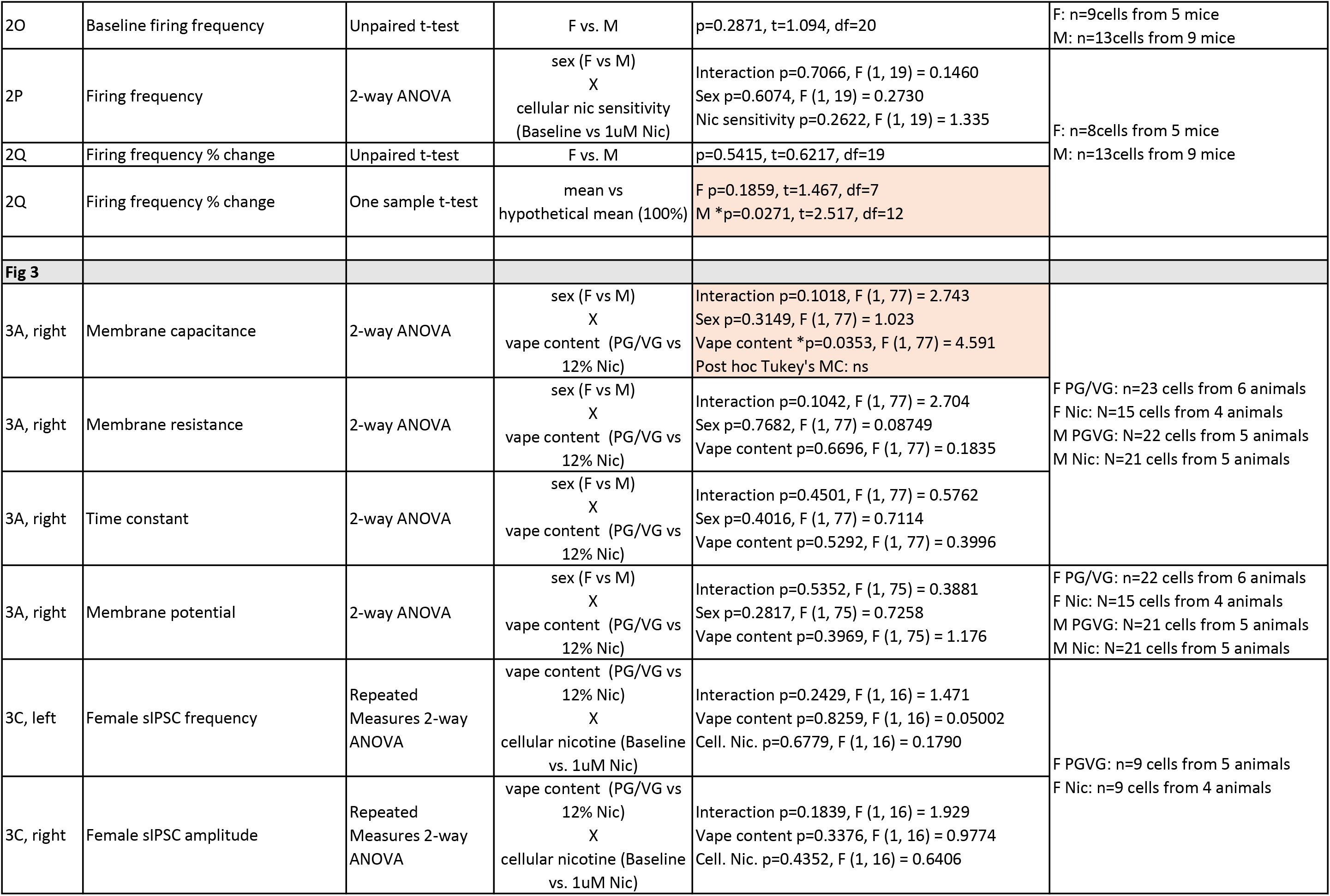
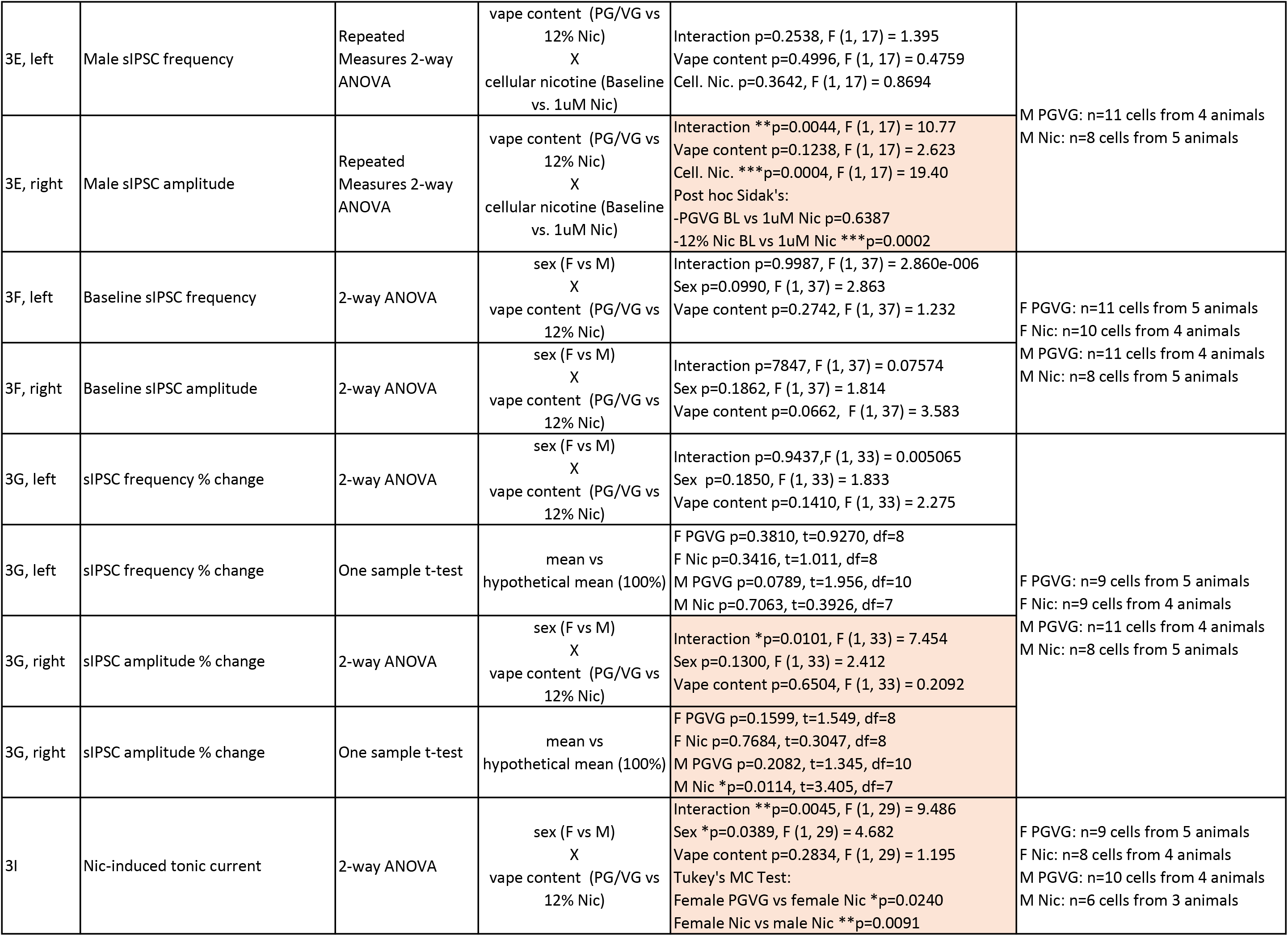
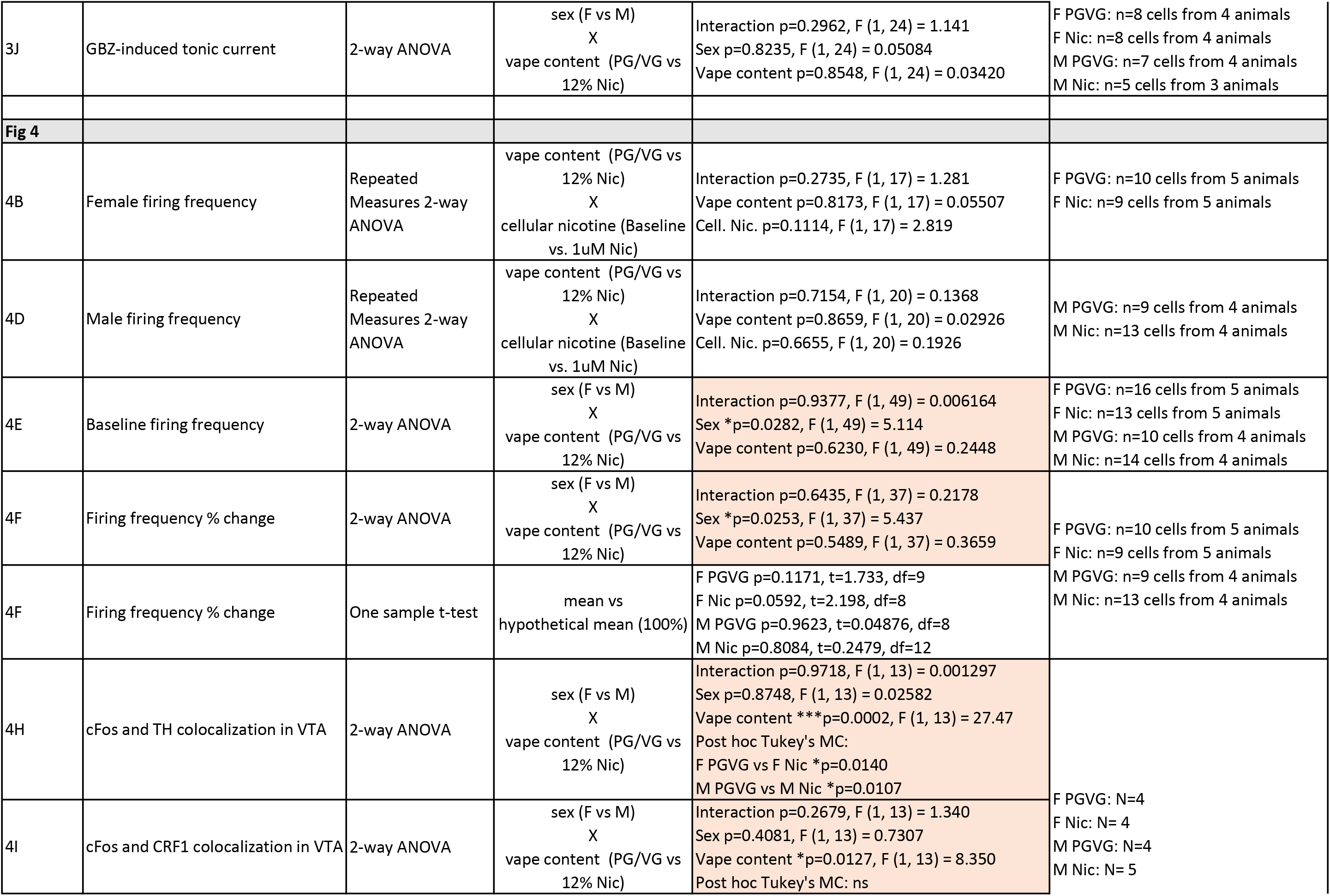
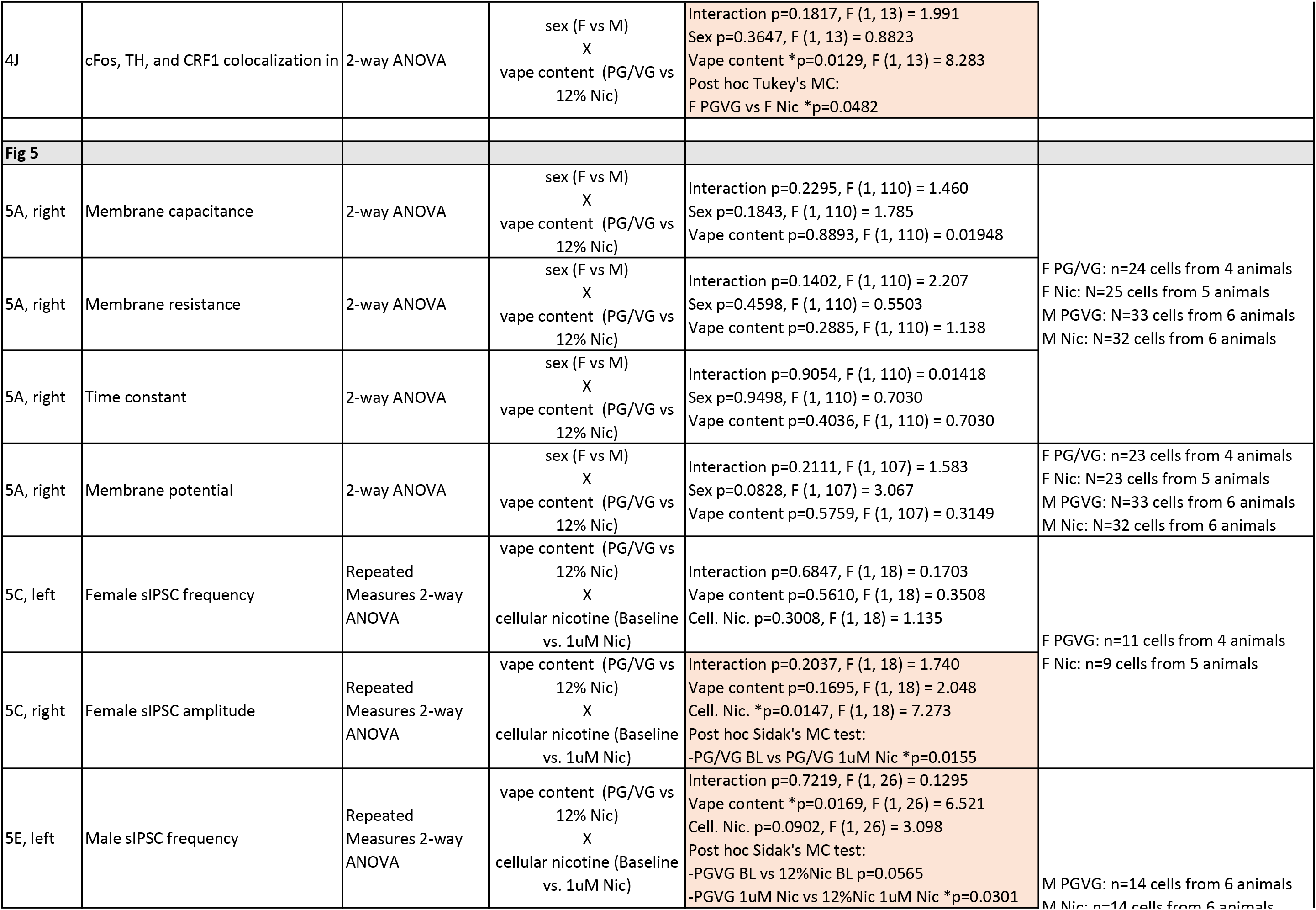
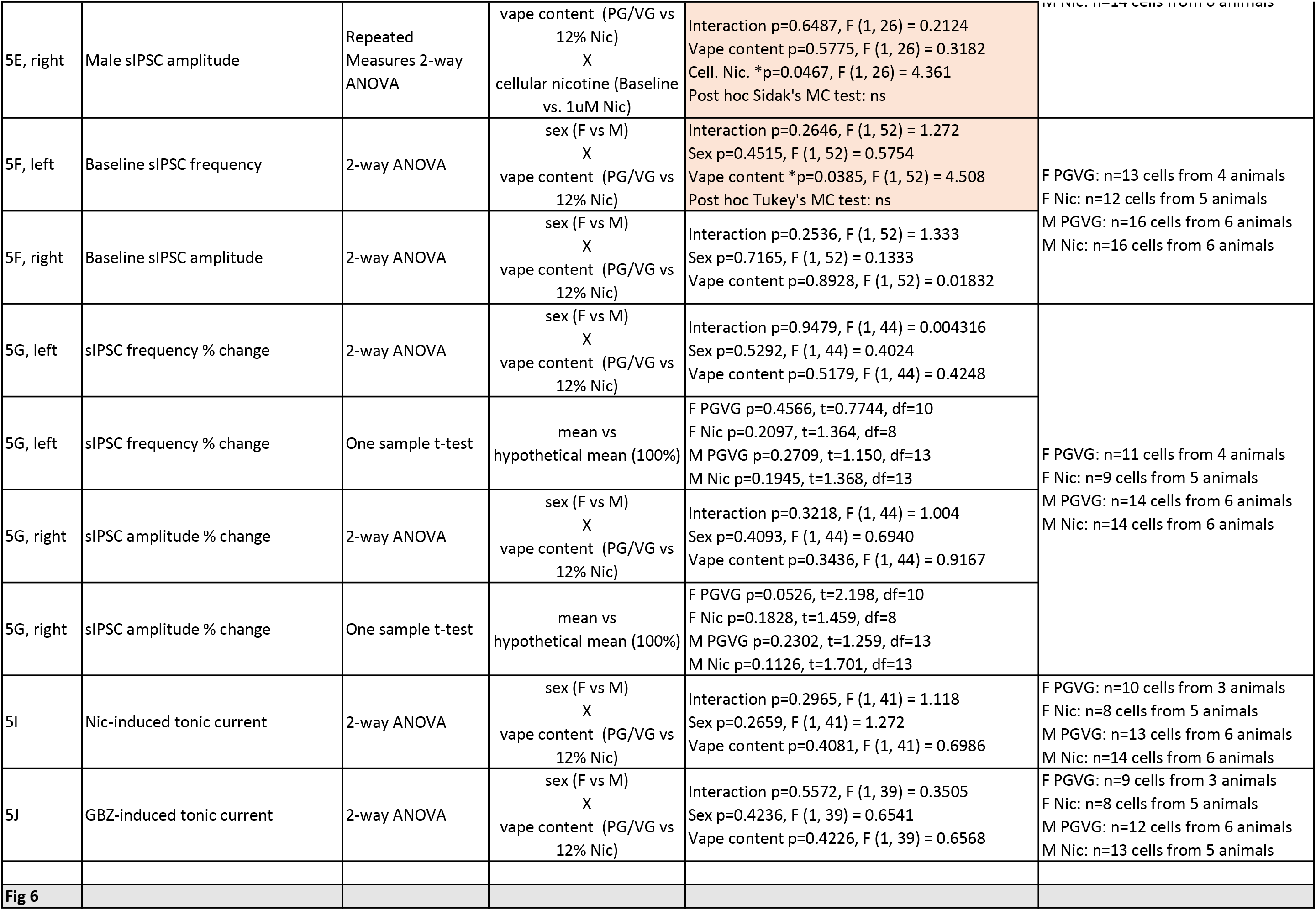
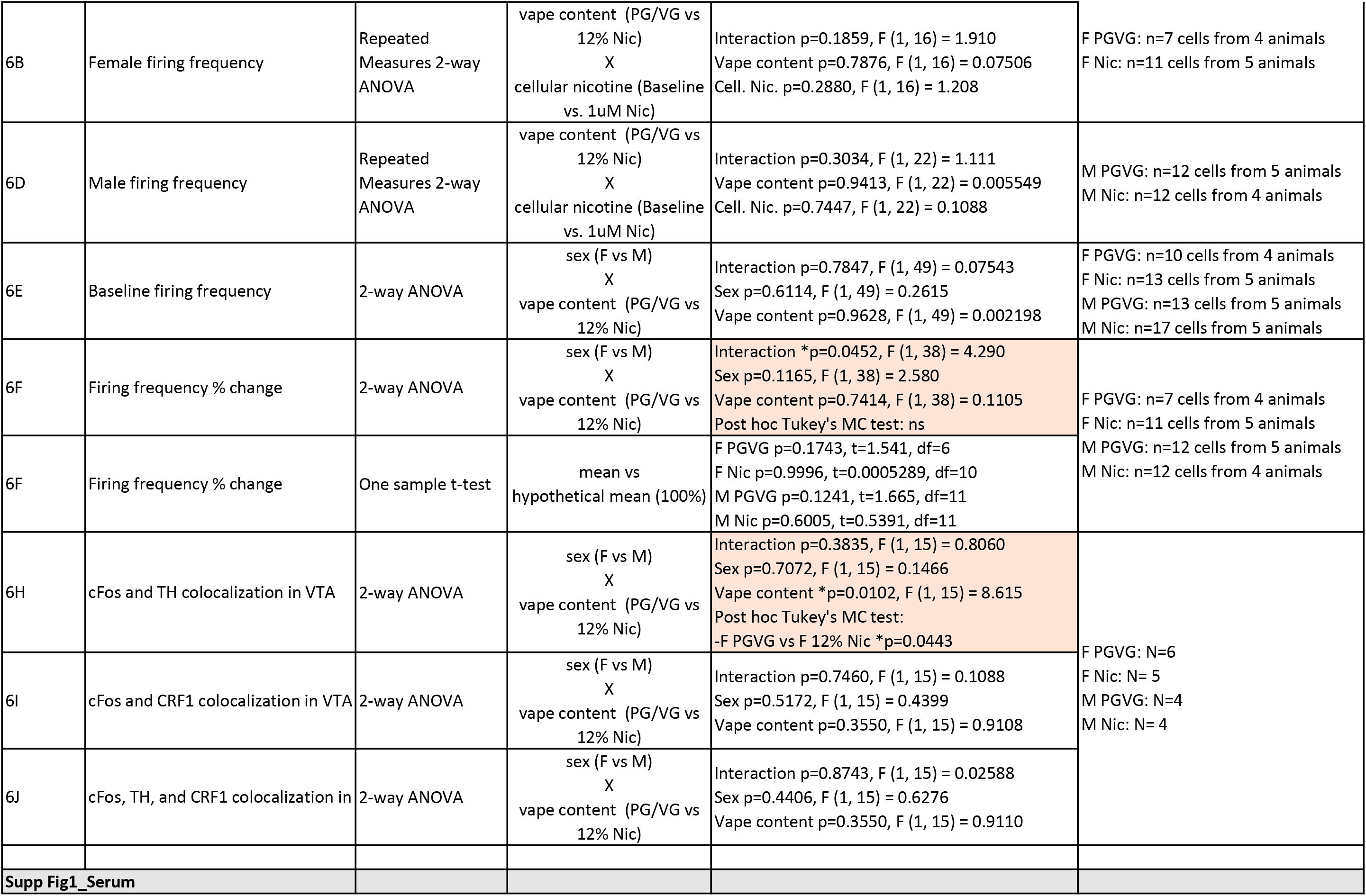
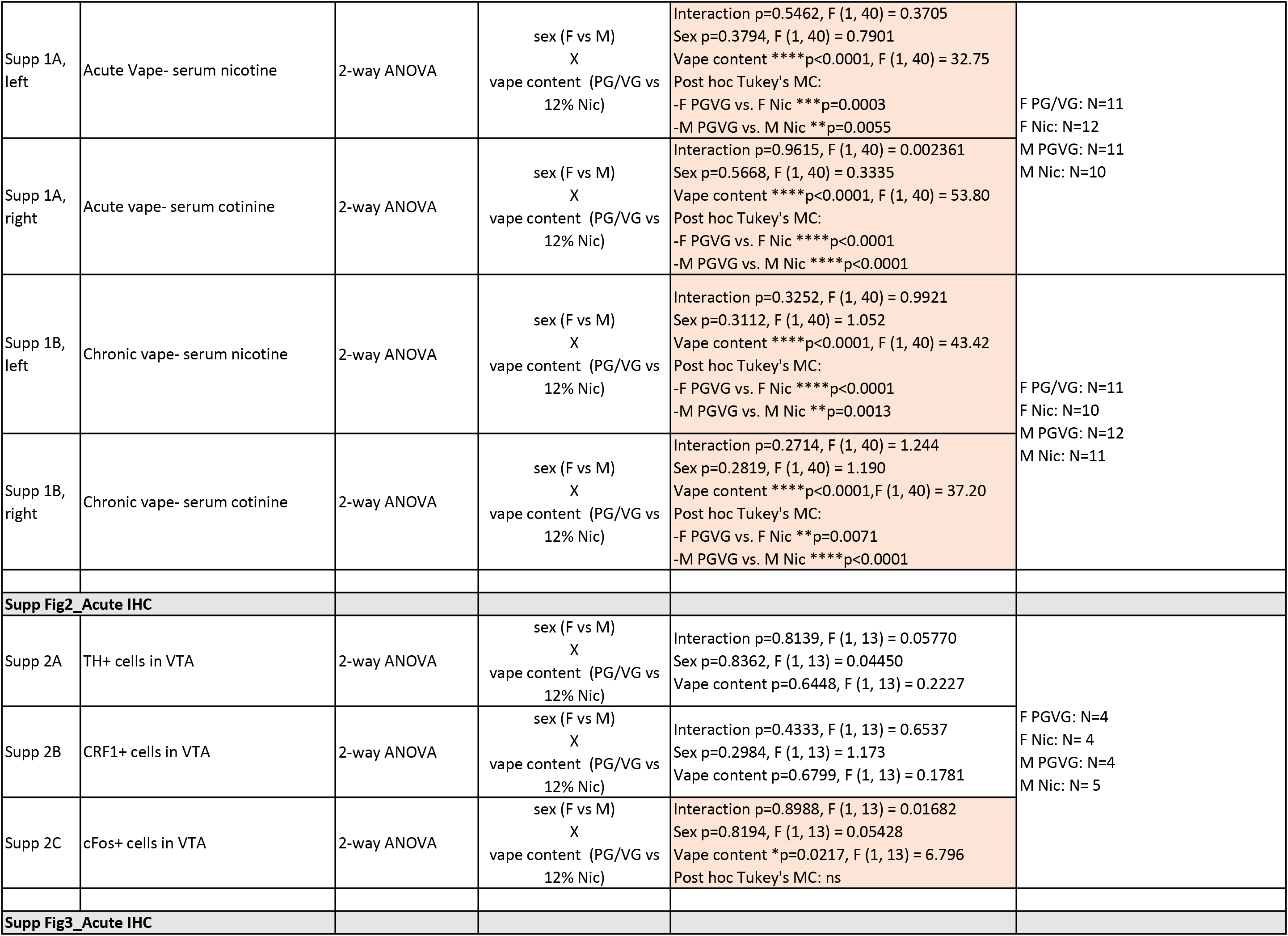
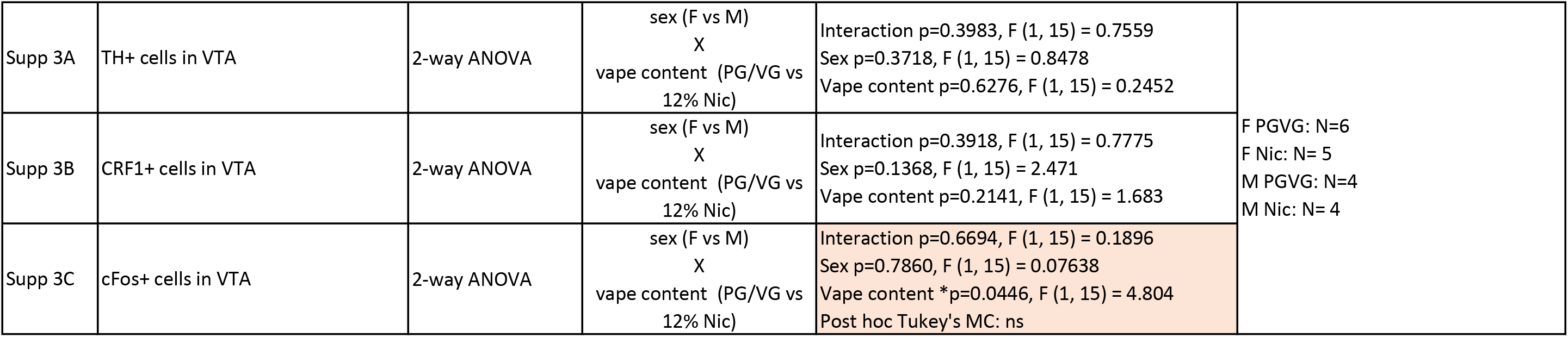

**Supplementary Figure 1:**
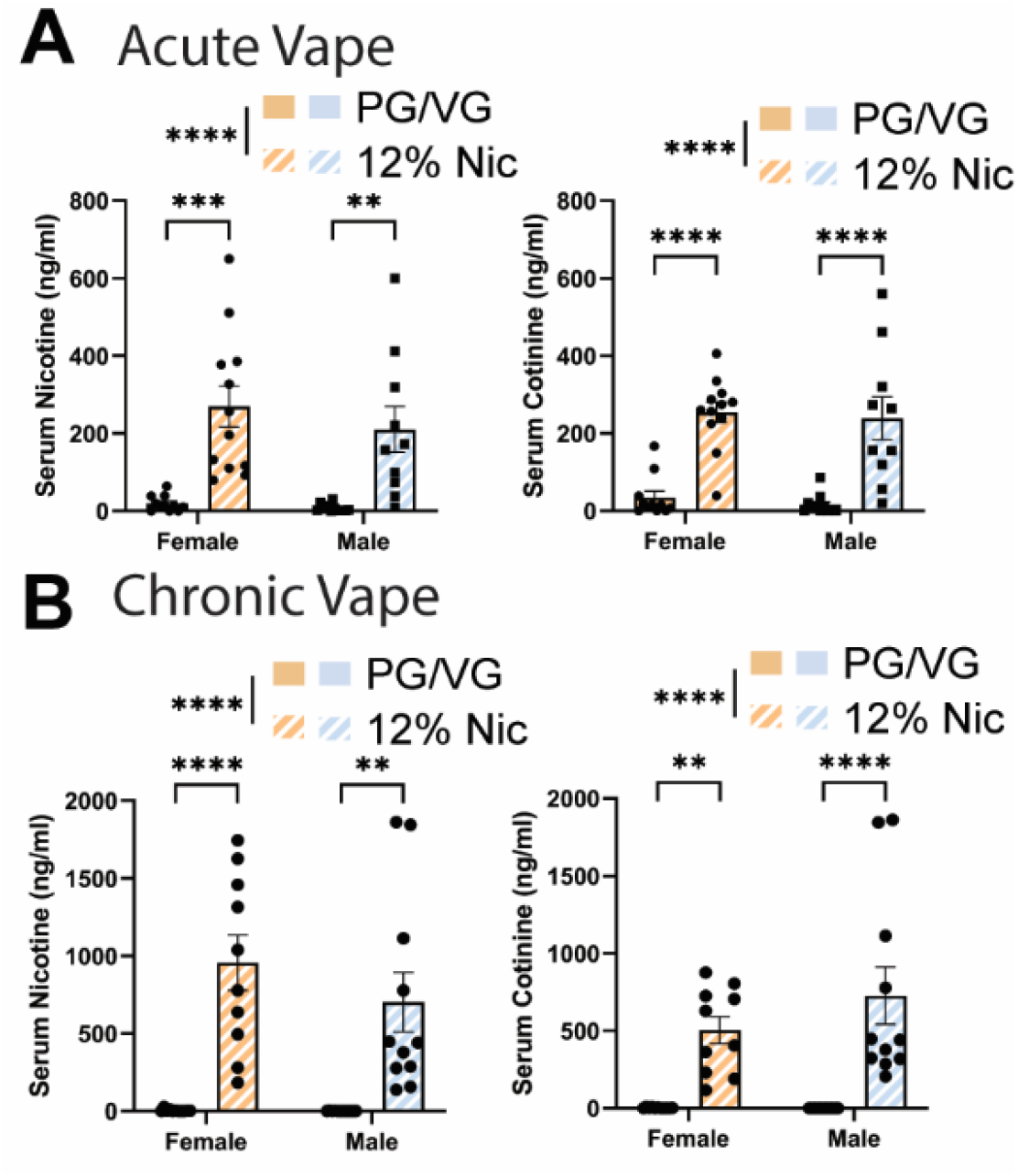
Nicotine and cotinine serum levels following acute and chronic electronic nicotine vapor exposure. **(A)** Serum nicotine (left) and serum cotinine (right) levels following acute PG/VG or 12% nicotine vape exposure in female and male mice. **(B)** Serum nicotine (left) and serum cotinine (right) levels following chronic PG/VG or 12% nicotine vape exposure in female and male mice. *\*\*p<0.005, ***p<0.0005, ****p<0.0001.* Error bars are SEM

**Supplementary Figure 2:**
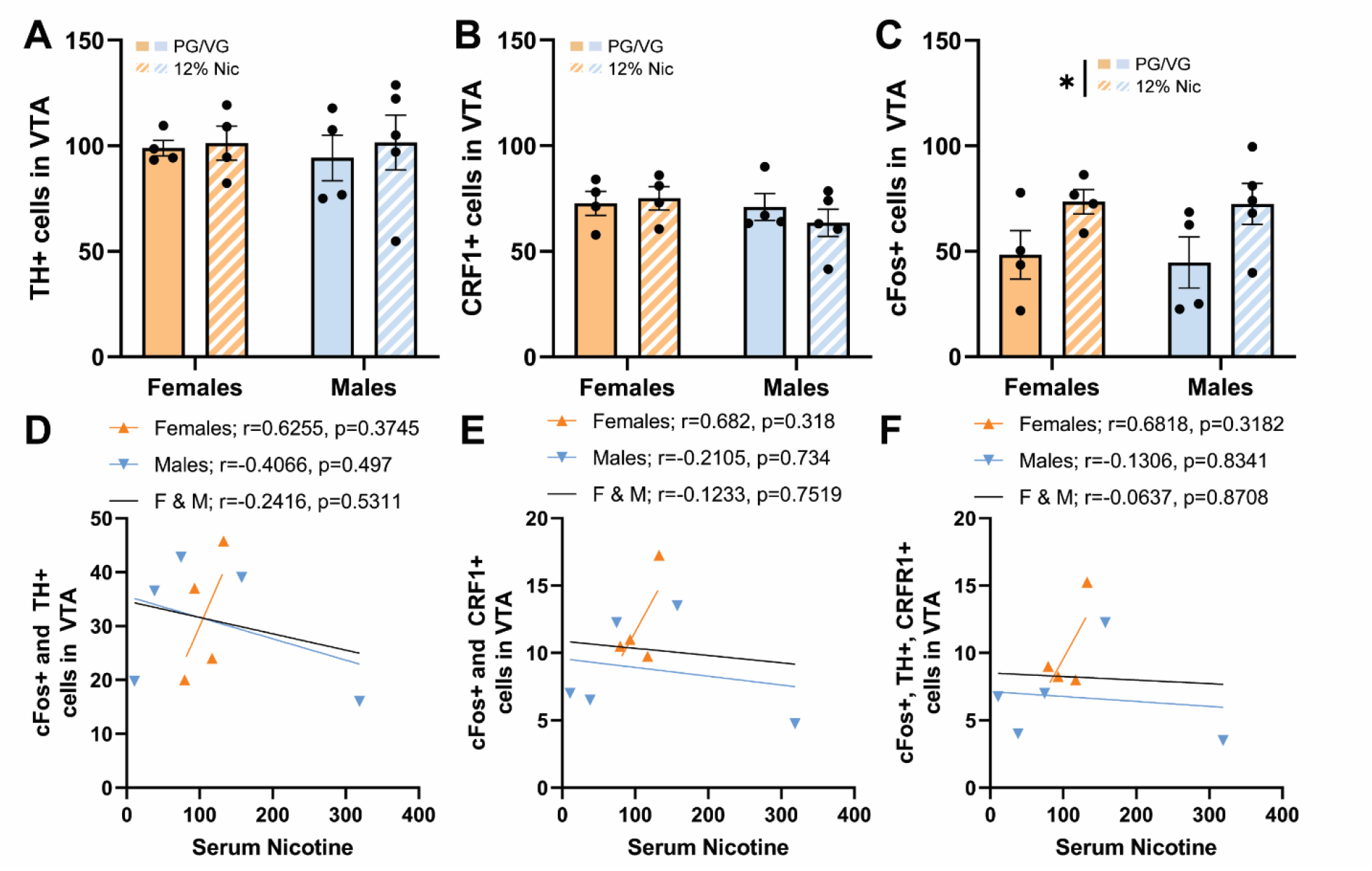
Acute vapor exposure immunohistochemistry and correlations with serum nicotine. **(A)** Number of TH+ neurons in the VTA in females and males exposed to acute PG/VG or acute 12% nicotine. **(B)** Number of CRF1+ neurons in the VTA in females and males exposed to acute PG/VG or acute 12% nicotine. **(C)** Number of cFos+ neurons in the VTA in females and males exposed to acute PG/VG or acute 12% nicotine. **(D)** Correlation between serum nicotine and number of cFos+ and TH+ cells in the VTA in females (orange), males (blue), and both sexes combined (black). **(E)** Correlation between serum nicotine and number of cFos+ and CRF1+ cells in the VTA in females (orange), males (blue), and both sexes combined (black). **(F)** Correlation between serum nicotine and number of cFos+, TH+, and CRF1+ cells in the VTA in females (orange), males (blue), and both sexes combined (black). *\*p<0.05.* Error bars are SEM

**Supplementary Figure 3:**
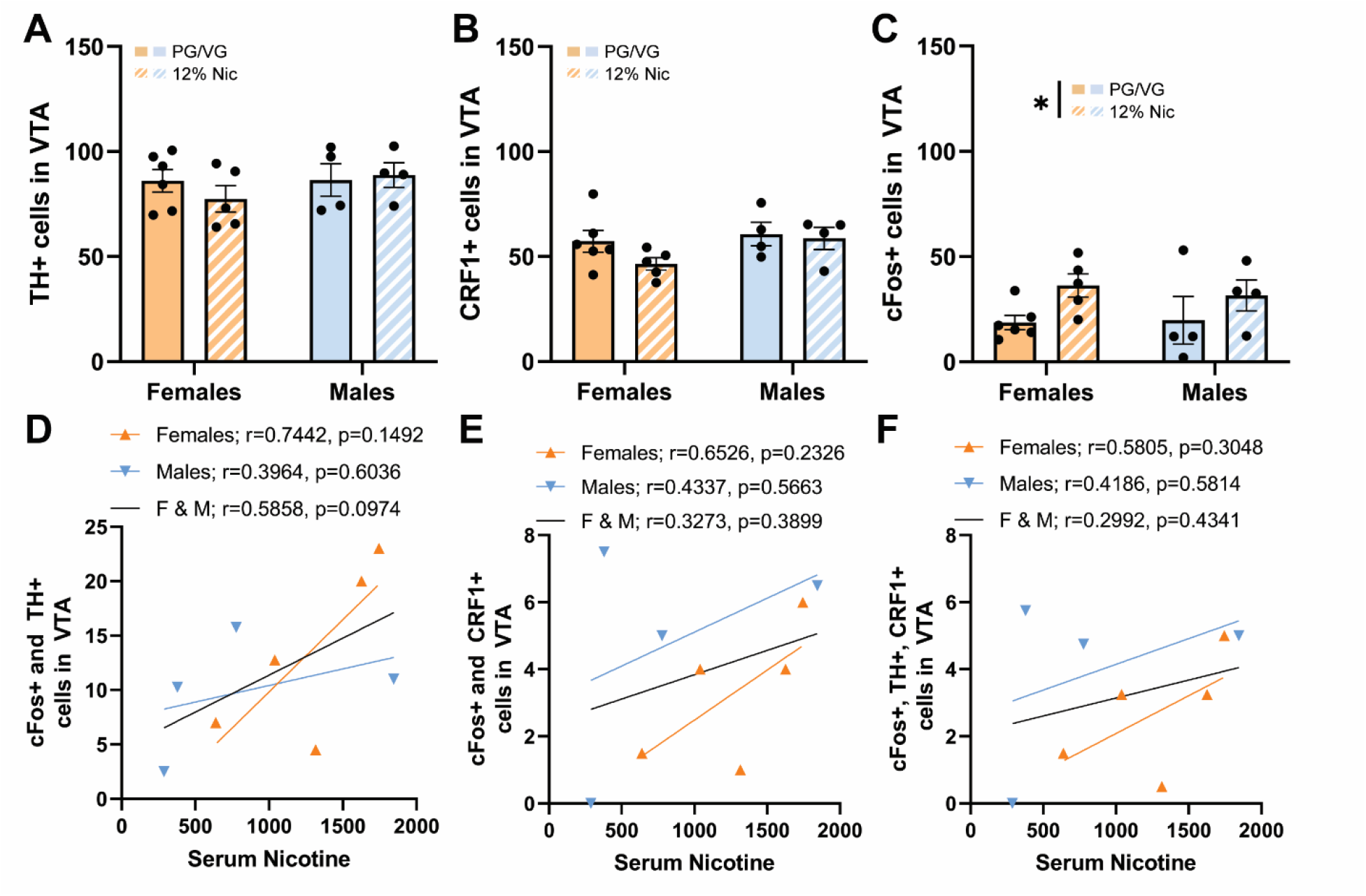
Chronic vapor exposure immunohistochemistry and correlations with serum nicotine. **(A)** Number of TH+ neurons in the VTA in females and males exposed to chronic PG/VG or chronic 12% nicotine. **(B)** Number of CRF1+ neurons in the VTA in females and males exposed to chronic PG/VG or chronic 12% nicotine. **(C)** Number of cFos+ neurons in the VTA in females and males exposed to chronic PG/VG or chronic 12% nicotine. **(D)** Correlation between serum nicotine and number of cFos+ and TH+ cells in the VTA in females (orange), males (blue), and both sexes combined (black). **(E)** Correlation between serum nicotine and number of cFos+ and CRF1+ cells in the VTA in females (orange), males (blue), and both sexes combined (black). **(F)** Correlation between serum nicotine and number of cFos+, TH+, and CRF1+ cells in the VTA in females (orange), males (blue), and both sexes combined (black). *\*p<0.05.* Error bars are SEM

